# Distinct spatial organization of human hippocampal perfusion and microvascular pulsatility across age

**DOI:** 10.64898/2026.10.15.759438

**Authors:** Fanhua Guo, Andy Jeesu Kim, Chenyang Zhao, Tanxin Dong, Zixuan Liu, Zidong Yang, Kay Jann, Xingfeng Shao, Neda Jahanshad, Mara Mather, Danny JJ Wang

**Affiliations:** Mark & Mary Stevens Neuroimaging and Informatics Institute, Keck School of Medicine, University of Southern California, Los Angeles, CA, USA; Department of Psychological Sciences, Purdue University, West Lafayette, IN, USA; Center on Aging and the Life Course, Purdue University, West Lafayette, IN, USA; Center for Affective and Autonomic Neuroscience, Max Planck Institute for Human Development, Berlin, Germany

## Abstract

Arterial pulsation can modulate neuronal activity through mechanosensitive pathways in mice, but whether vascular pulsatility is similarly related to neural organization in humans remains unclear. Here, using 7T MRI in 35 young and older adults, we mapped cerebral perfusion and microvascular volumetric pulsatility index (mvPI) across the unfolded hippocampus and integrated these measures with histological, electrophysiological, and transcriptomic reference maps. Perfusion and mvPI exhibited distinct spatial organization and age-related differences. In young adults, perfusion preferentially corresponded to parvalbumin-interneuron density and beta/gamma distribution, whereas mvPI was associated with theta-band distribution. A number of associations detected in young adults were not detected in the older adults maps. Spectral simulations showed qualitative similarity to heartbeat-related phase resetting as one possible explanation for the mvPI–theta relationship, while spatial transcriptomics revealed perivascular enrichment of *PIEZO2*, a candidate mediator of vascular mechanotransduction. Together, these findings establish perfusion and tissue-level pulsatility as distinct dimensions of human hippocampal vascular organization and provide a framework for investigating their relationships with neural dynamics across aging.

## Introduction

The brain has negligible energy reserves, so neuronal activity depends on the continuous delivery of oxygen and nutrients by the cerebral vasculature^1,2^. The vasculature has therefore been viewed chiefly as a metabolic supply system matched to local demand through neurovascular coupling^3,4^. Beyond this metabolic role, however, the cerebral vasculature is also intrinsically pulsatile. Each heartbeat propagates a pressure wave through the arterial tree^5^, producing rhythmic changes in vascular volume and mechanical forces within the surrounding tissue^6^. The vascular environment thus has two distinct dimensions: a mean-flow component that determines tissue perfusion and a pulsatile component that imposes rhythmic mechanical forces on the parenchyma. Whereas the former is central to conventional models of neurovascular coupling, the latter has been studied primarily as a feature of vascular physiology and pathology^7,8^, with much less attention to its potential role as a direct modulator of neural activity.

Several lines of evidence suggest that this view is incomplete. Arterial pulsation drives perivascular cerebrospinal fluid transport and solute clearance^9–11^, indicating physiological roles beyond blood delivery. Neurons can also respond directly to mechanical stimuli: mechanically gated cation channels can trigger action potentials in neocortical and hippocampal neurons^12^, and the mechanosensitive channel Piezo2 is expressed in select neuronal populations^13^. Most directly, Salameh et al.^14^ showed that blood pressure pulsations modulate neuronal activity through Piezo2 in mice, establishing a vascular-to-neural signaling pathway opposite to classical neurovascular coupling. In humans, cardiac-cycle-locked neural activity has been detected with intracranial recordings^15–18^, EEG^19,20^ and MEG^21^, with prominent effects in the theta range^15,21^. However, whether these responses arise from interoceptive and autonomic pathways or, at least partly, from local mechanical transduction of vascular pulsation remains unresolved. Distinguishing these mechanisms in humans has been challenging because vascular mechanical inputs cannot be directly manipulated, while conventional measurements provide limited ability to spatially resolve pulsatility within brain tissue.

The hippocampus provides a favorable setting to examine whether perfusion and pulsatility show distinct relationships with neural organization. Its arterial supply consists of branches of the anterior choroidal and posterior cerebral arteries enter predominantly from the inner surface and follow distinct courses along the hippocampal long axis^22–25^. This vascular architecture may impose distinct spatial constraints on the delivery of blood flow and arterial pulsations, providing an anatomical basis for local perfusion and pulsatile load to exhibit partially dissociable spatial organization. The hippocampus also exhibits neural features potentially relevant to these vascular properties. It is characterized by prominent theta activity^26^, and heartbeat-related neuronal activity has been observed in the mouse hippocampus^14^ (with Piezo2 neurons^12–14^) and in human intracranial recordings from hippocampal regions^16–18^. In parallel, the hippocampus contains a prominent population of fast-spiking parvalbumin-positive (PV+) interneurons^27–29^ that support high-frequency activity at substantial metabolic cost, providing candidate neural features related to pulsatile and metabolic aspects of the vascular environment, respectively. These vascular and neural properties are embedded within continuous functional, structural and molecular gradients, particularly along the anterior–posterior axis^30–35^. HippUnfold^36^ and HippoMaps^37^ toolkits now enable these features to be compared within a common surface-based coordinate system. Finally, the hippocampus exhibits pronounced vulnerability to age-related and vascular alterations^38–40^, providing an opportunity to examine whether these vascular–neural relationships also diverge with age.

Aging may differentially affect these two vascular components. Stiffening of central elastic arteries reduces pulse buffering and increases pulsatile energy transmitted to the cerebral circulation^41^, while cerebral perfusion declines with age^42^. In the hippocampus, vascular supply has been associated with regional volume and cognitive performance^38,43^. Yet perfusion and pulsatility have rarely been examined together at the tissue level. Pulsatility is typically measured in large arteries using transcranial Doppler^44^ or phase-contrast MRI^45^, providing limited information about its distribution within hippocampal tissue. It therefore remains unclear whether tissue perfusion and pulsatility exhibit distinct spatial organization, whether they show distinct relationships with neural organization, and whether these relationships diverge with age.

Recent methodological advances provide an opportunity to address these questions in the human hippocampus. First, advances in ultrahigh-field vascular MRI now enable both microvascular perfusion and pulsatility to be resolved at the tissue level. With 7T MRI, a FLASH-based rotated golden-angle stack-of-spirals (rGA-SoS) pseudocontinuous ASL (pCASL) sequence provides whole-brain perfusion mapping at 1-mm isotropic resolution^46,47^, while 4D vascular space occupancy (VASO) MRI^48,49^ enables quantification of microvascular volumetric pulsatility index^48^ (mvPI), capturing cardiac-related microvascular volume changes within tissue. And recent implementations have further demonstrated the feasibility of high-resolution VASO imaging in the hippocampus^50^. Second, surface-based hippocampal frameworks overcome the complex folded geometry of this structure: HippUnfold provides subject-specific unfolding and subfield segmentation^36^, while HippoMaps brings cytoarchitectonic, electrophysiological and functional reference maps into a common coordinate system^37^. Finally, the latest human hippocampal spatial transcriptomic atlases^51^ provide complementary molecular information at cellular-scale spatial resolution. Together, these advances make it possible to relate regional perfusion and pulsatility to neural organization across imaging, electrophysiological and molecular scales within the human hippocampus.

Here, we investigated whether cerebral perfusion and vascular pulsatility exhibit distinct spatial organization in the human hippocampus, whether they show differential relationships with neural architecture, and how these patterns differ with age. We acquired multimodal 7T MRI in 15 young and 20 older adults, combining high-resolution CBF and mvPI measurements with time-of-flight angiography (TOF-MRA) to characterize perfusion, tissue-level pulsatility, and arterial anatomy within the same individuals. We first examined the spatial organization of CBF and mvPI across the hippocampus and their differences between age groups. We hypothesized that CBF, as an index of metabolic substrate delivery, would preferentially correspond to neural features associated with high energetic demand, whereas mvPI would show a distinct pattern of association with hippocampal neural organization. We then examined whether these vascular–neural relationships differed with age by integrating the vascular measurements with reference maps of cytoarchitecture, intracranial electrophysiology, and resting-state functional organization. Finally, human hippocampal spatial transcriptomics was used to examine the spatial organization of candidate molecular substrates of vascular mechanotransduction. Together, this multimodal framework allowed us to characterize distinct relationships of perfusion and pulsatility with hippocampal organization across vascular, neural, and molecular scales.

## Results

### Hippocampal Subfield Differences in Perfusion and Pulsatility with Age

We first measured cerebral blood flow (CBF) with iso-1mm resolution in each participant using our rGA-SoS pCASL sequence at 7T^46^. Microvascular volumetric pulsatility index^48^ (mvPI) was quantified with iso-1.2mm resolution using dynamic VASO MRI scans^49^ at 7T (see Methods for details). High-resolution structural images (iso-0.7mm) were then segmented using the HippUnfold toolkit^36^. **Fig. 1a** shows the resulting segmentation of the hippocampus and its subfields in a representative participant. **Fig. 1b** presents the mean CBF across hippocampal subfields in the young and older groups respectively. An age (2) × subfield (7) mixed-design repeated-measures ANOVA revealed significant main effects of age (F(1,30) = 8.71, Greenhouse-Geisser corrected p = 6.1×10^-3^, partial η^2^ = 0.23) and subfields (F(6,180) = 10.62, GG-corrected p = 4.0×10^-7^, partial η^2^ = 0.26), with no significant age × subfield interaction (F(6,180) = 1.64). Post-hoc t-tests showed significant age-related differences in all subfields except CA3 (BH-FDR corrected). These findings indicate significant regional differences in CBF across hippocampal subfields. Moreover, hippocampal CBF was lower in older adults, with no evidence that the magnitude of this age-group difference varied across subfields. **Fig. 1c** presents the mean mvPI across hippocampal subfields in the young and older groups. A 2 (age) × 7 (subfields) mixed-design repeated-measures ANOVA showed significant main effects of age (F(1,30) = 10.09, GG-corrected p = 3.4×10^-3^, partial η^2^ = 0.25) and subfields (F(6,180) = 16.43, GG-corrected p = 4.4×10^-9^, partial η^2^ = 0.35), as well as a significant age × subfield interaction (F(6,180) = 5.82, GG-corrected p = 8.0×10^-4^, partial η^2^ = 0.16). Post hoc t-tests identified significant age-related differences in the subiculum (Sub) and CA1-4 (BH-FDR corrected). These results demonstrate significant regional differences in mvPI across hippocampal subfields. Furthermore, hippocampal mvPI increased with age, with the magnitude of the increase varying across subfields. The normalized ΔVASO pattern closely resembled that of mvPI (**Fig. S1a**). Across participants, variance decomposition further showed that ΔVASO accounted for substantially greater unique variance in mvPI than CBF across all hippocampal subfields (**Fig. S1b**), indicating that interindividual variation in mvPI was predominantly driven by its VASO-derived pulsatile component. Together, these findings demonstrate distinct spatial patterns of CBF and mvPI across hippocampal subfields. While CBF exhibited a uniform age-related decline across subfields, age-related increases in mvPI were subfield dependent.

**Fig. 1.**
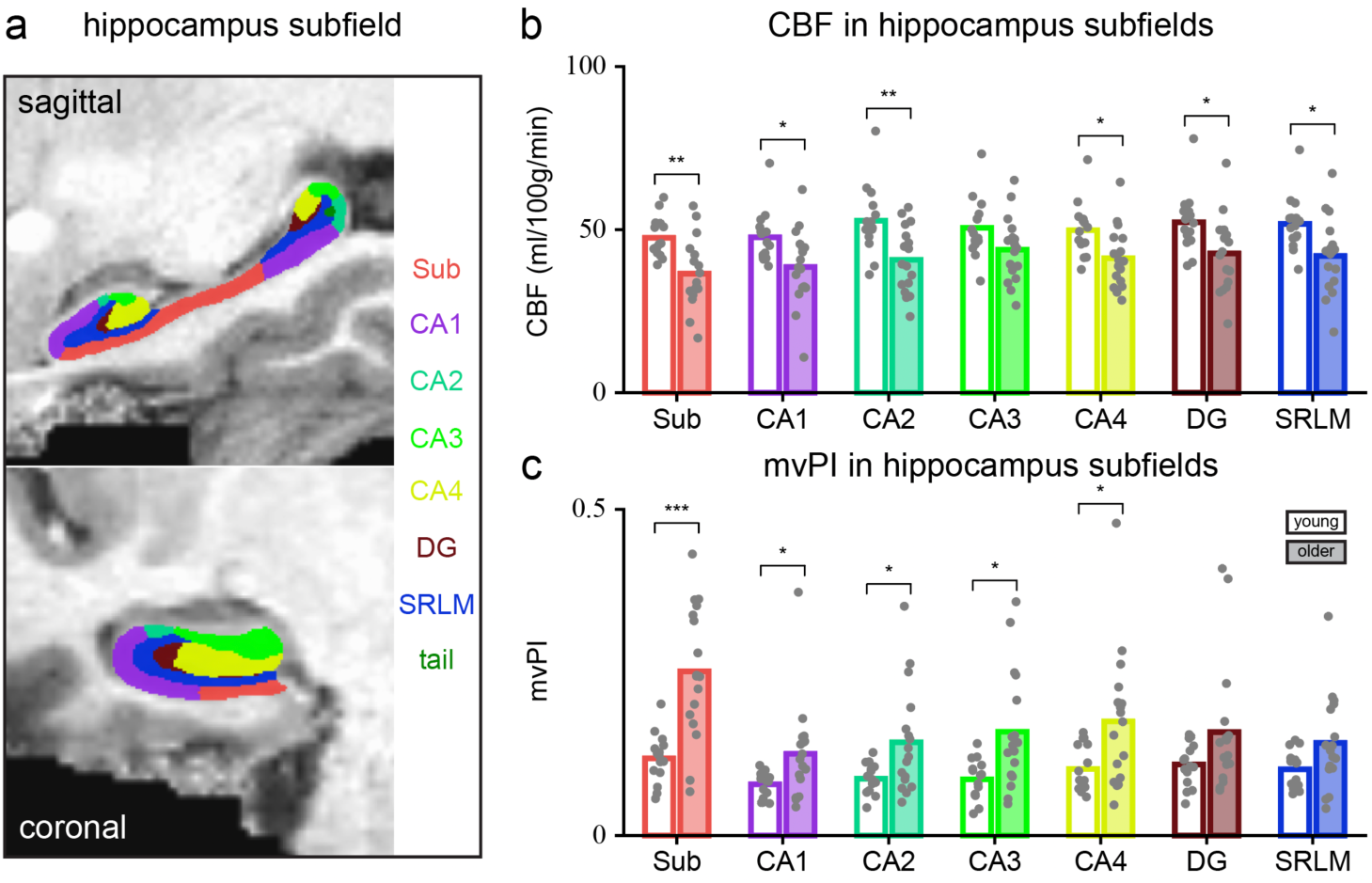
CBF and mvPI show distinct subfield organization and distinct age differences in the human hippocampus. **a**, Hippocampal segmentation in a representative participant. High-resolution T1-weighted images (0.7-mm isotropic) were segmented with HippUnfold into seven subfields: subiculum (Sub), cornu ammonis (CA) 1, CA2, CA3, CA4, dentate gyrus (DG) and stratum radiatum, lacunosum, and moleculare (SRLM) and colour-coded as in **b**,**c**. **b**, Mean CBF in each subfield in young (n = 15) and older (n = 17) adults. Bars give the group mean across participants and dots show individual participants. Main effects of age and subfield and their interaction were assessed with a 2 (age) × 7 (subfield) mixed-design repeated-measures ANOVA (Greenhouse–Geisser corrected). Horizontal lines and asterisks denote post hoc between-age comparisons within each subfield (2-sided unpaired T-test, BH-FDR corrected across subfields). Sub/CA1/CA2/CA3/CA4/DG/SRLM FDR = 0.009/0.019/0.009/0.058/0.019/0.019/0.019. **c**, As in **b** for mvPI. Sub/CA1/CA2/CA3/CA4/DG/SRLM FDR = 0.0003/0.037/0.027/0.019/0.027/0.075/0.059. Asterisks in **b**,**c**: *p < 0.05, **p < 0.01, ***p < 0.001, BH-FDR corrected.

### Spatial Distribution of Hippocampal Perfusion and Pulsatility with Age

Beyond its distinct subfield organization, the hippocampus exhibits continuous functional^30–33^, structural^33,34^, and molecular^33,35^ gradients across its unfolded surface, particularly along the anterior-posterior (AP) axis. Aging has also been shown to exert spatially heterogeneous effects on hippocampal function^31^ and structure^52–54^. We therefore performed a spatially resolved analysis of hippocampal CBF and mvPI using the HippUnfold^36^ and Hippomaps^37^ frameworks.

**Fig. 2a** shows the average unfolded hippocampal CBF maps for the young and older groups. In young group, high CBF was observed across most of the CA2/3 regions and the posterior hippocampus. In contrast, older group exhibited relatively elevated CBF primarily within the middle CA2/3, where it appeared as spatially separated patches. **Fig. 2b** presents the corresponding unfolded mvPI maps. In both groups, mvPI was higher in the anterior hippocampus and middle subiculum. **Fig. S2a** shows the spatial distribution of normalized ΔVASO across the unfolded hippocampus. The ΔVASO pattern closely resembled that of mvPI, suggesting that although mvPI is derived from both ΔVASO and CBF, its spatial variation is predominantly driven by ΔVASO, consistent within subfields. **Fig. 2c** and **Fig. S2b** show the probability density distribution (PDD) maps of large arteries (about diameter ≥ 0.5 mm) located adjacent to the hippocampal inner and outer surface, respectively, derived from TOF images with iso-0.5mm resolution. The PDD map of arteries adjacent to the hippocampal inner surface closely matched the mvPI distribution, suggesting that hippocampal arterial anatomy contributes to the spatial organization of mvPI. This observation is consistent with previous reports that elevated cortical surface mvPI is primarily driven by pial arteries^48^. Furthermore, excluding TOF-visible arteries and adjacent vertices minimally affected the mvPI spatial pattern (young: r = 0.97; older: r = 0.98; **Fig. S2e**), indicating that the pattern was robust to potential partial-volume effects from large arteries. **Fig. 2d** presents the spatial correlation matrix of these maps, calculated using Pearson correlation with the spin-null model to correct for spatial autocorrelation (BH–FDR corrected). Inner surface arterial distribution, mvPI, and ΔVASO exhibited strong positive spatial correlations with one another (r > 0.8, FDR < 0.001). Next, we examined the spatial distribution of age-related changes in CBF and mvPI. **Fig. S2c** and **Fig. 2e** show the difference and normalized difference maps of CBF between the young and older groups, together with clusters exhibiting significant group differences (permutation test, vertex-wise p < 0.001). Significant age-related CBF decline was concentrated primarily in the posterior hippocampus, corresponding largely to the vascular territory of the posterior hippocampal artery^43^. **Fig. S2d** and **Fig. 2f** present the corresponding mvPI maps and significant clusters (permutation test, vertex-wise p < 0.001, BH–FDR corrected). Significant age-related increases in mvPI were distributed across three regions: the anterior hippocampus, the posterior hippocampus, and the middle subiculum. **Fig. 2g** shows the changes in CBF and mvPI profiles across the four subfields along the anterior-posterior axis and the age-related changes are consistent with the maps. Since posterior hippocampal atrophy has been reported during normal aging^52–54^, we examined regional hippocampal volume (anterior, middle and posterior) as a function of age. Regression models including linear and quadratic age terms, with sex and estimated total intracranial volume (eTIV) as covariates, revealed significant age-related volume reductions in the middle and posterior hippocampus, but not in the anterior hippocampus (**Fig. 2h**). This pattern only partially overlapped with the regional perfusion changes: the middle hippocampus showed volume reduction without a corresponding significant decline in CBF, whereas the posterior hippocampus showed both volume and CBF reductions. Consistent with this spatial convergence, regional volume was positively associated with CBF in the posterior hippocampus after accounting for eTIV (**Fig. 2i**), whereas no significant associations were observed in the anterior or middle segments. These findings indicate partially overlapping spatial patterns of structural and perfusion-related age differences along the hippocampal long axis, with the clearest convergence in the posterior hippocampus.

**Fig. 2.**
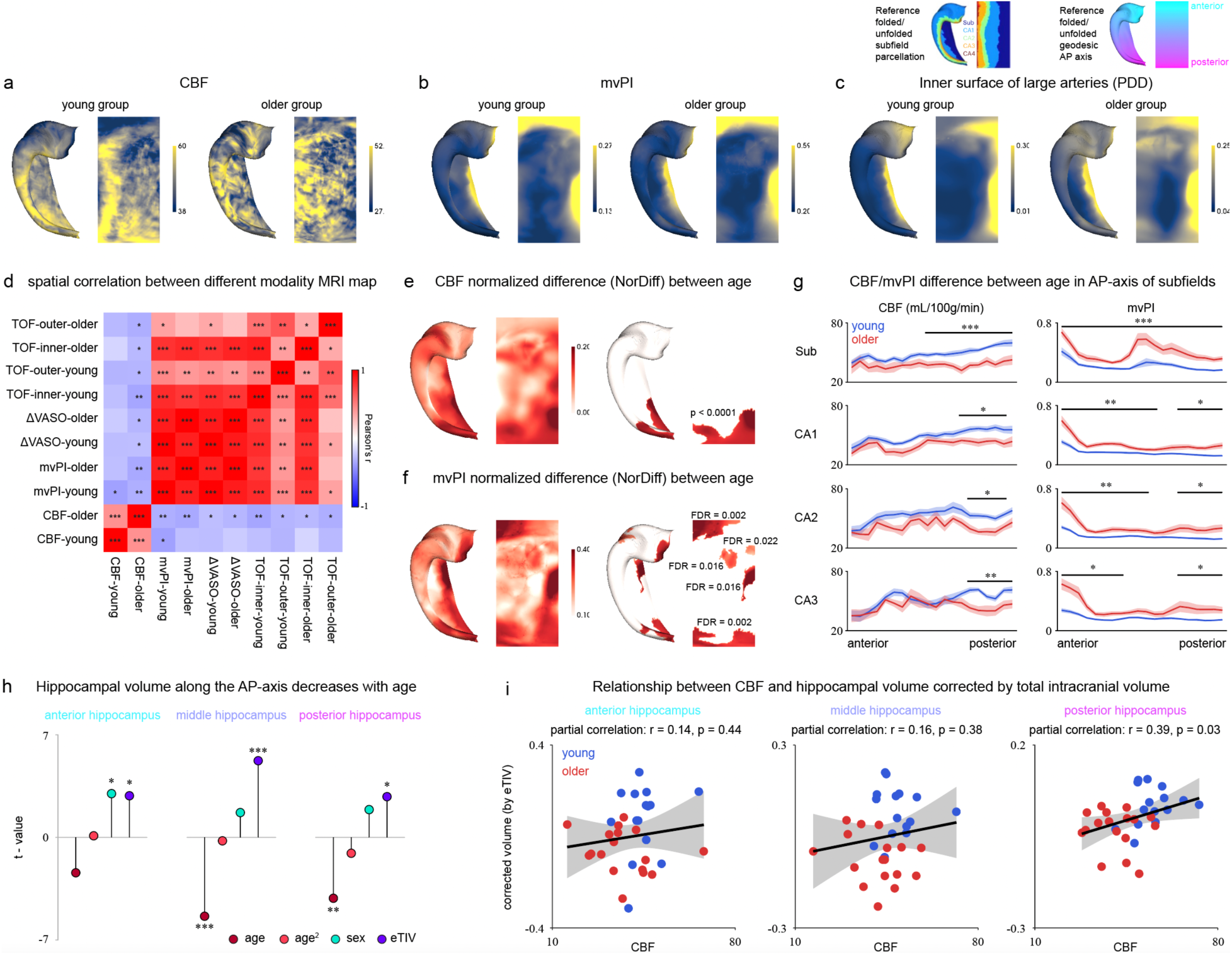
CBF and mvPI unfolded hippocampal spatial maps and age differences. **a**, Group-average CBF maps in unfolded hippocampal space for young (left, n = 15) and older (right, n = 17) adults. Color bar gives CBF in ml/100 g/min. **b**, Group-average mvPI maps in unfolded hippocampal space for young (left, n = 15) and older (right, n = 17) adults. **c**, Probability density distribution (PDD) of large arteries (diameter ≥ 0.5 mm) adjacent to the inner hippocampal surface, derived from TOF angiograms and averaged across young (left, n = 15) and older (right, n = 16) adults. Values give the across-participant probability of a large artery being present within each vertex. **d**, Spatial correlation matrix among the maps in (**a–c**) and **Fig. S2**. Entries give vertex-wise Pearson correlations and significance p value was assessed against null distributions obtained from 10000 spatial surrogate maps generated with spin-null model, which preserve the spatial autocorrelation of each map. Asterisks: *p < 0.05, **p < 0.01, ***p < 0.001, BH-FDR corrected across the matrix. **e**, Normalized age difference in CBF, computed vertex-wise as (young − older)/(young + older). Right panel shows the significant clusters of group differences (permutation cluster test, vertex-wise p < 0.001, BH-FDR correction). **f**, As in **e** for mvPI. **g**, CBF (left) and mvPI (right) along the AP axis, shown separately for Sub, CA1, CA2 and CA3 in young (blue) and older (red) adults. The AP axis was divided into 20 equally spaced bins from anterior to posterior. Lines and shaded bands give the mean and s.e.m. across participants. Horizontal bars denote bins with significant between-age differences (permutation cluster test, BH-FDR corrected). **h**, T-statistics for the associations of age with hippocampal volume in the anterior, middle and posterior segments. Regional hippocampal volume was modeled as a function of age and age², with sex and estimated total intracranial volume (eTIV) included as covariates. **i**, Associations between regional hippocampal volume and CBF in the anterior (left), middle (center) and posterior (right) hippocampus. Hippocampal volumes were adjusted for eTIV before correlation analysis. Each point represents one participant, with young and older adults shown in blue and red, respectively. Black lines indicate linear regression fits and shaded areas indicate 95% confidence intervals. Values indicate partial correlation coefficients (r) and BH-FDR corrected p values across the three hippocampal segments.

### Spatial Relationships between Vascular Features and Hippocampal Structure and Function

The brain maintains minimal endogenous energy reserves, with glycogen stored in limited quantities and largely confined to astrocytes^1,2^. Neuronal activity therefore depends on the continuous delivery of metabolic substrates by the cerebral vasculature, suggesting that the cellular architecture and functional organization of the hippocampus may be spatially coupled to its vascular organization. Hippocampal arterial pulsation has additionally been reported to modulate neuronal activity directly^14^. We therefore examined the spatial correspondence between our cerebrovascular feature maps (CBF and mvPI) and multimodal hippocampal reference maps spanning cell-stain distributions, resting-state fMRI (rs-fMRI) and intracranial EEG (iEEG).

**Fig. 3a,b** shows the reference maps: five cell stains (Merker, Calbindin, Calretinin, Parvalbumin, Thionin), three rsfMRI metrics (intrinsic timescale, IntTS; regional homogeneity, reho; functional connectivity with cortex, FC) and iEEG power in five frequency bands (delta, theta, alpha, beta, gamma). Partial least squares (PLS) regression was performed separately in the young and older groups to identify latent components linking cerebrovascular features to hippocampal structure-function organization. The first latent component (LC1) accounted for 35.7% and 29.2% of the cross-block covariance between cerebrovascular features and hippocampal reference maps in young and older groups, respectively. In both groups, LC1 (**Fig. 3c**) was defined by an opposition between the two vascular measures, with CBF at its negative pole (r = -0.91 young, -0.75 older) and mvPI at its positive pole (r = 0.73 and 0.88, respectively). Negative LC1 loadings therefore denote maps covarying positively with CBF, and positive loadings maps covarying positively with mvPI. The relative weighting of the two measures shifted modestly between groups, from slightly CBF-dominated in the young group to slightly mvPI-dominated in the older group. Replacing mvPI with its CBF-independent ΔVASO component yielded the same opposing pattern (**Fig. S3**), indicating that it was not driven by mathematical dependence on CBF. In the young group, LC1 loadings exceeding spatial-autocorrelation-preserving null distributions were restricted to Parvalbumin-stain and to theta, beta and gamma power (spin-null p < 0.05, BH-FDR corrected). Beta and gamma loaded at the CBF pole and theta at the opposite pole, and the underlying vertex-wise correlations (**Fig. 3d,e**) supported a frequency-specific relationship in which CBF was positively aligned with high-frequency power (beta r = 0.77, gamma r = 0.73; spin-null FDR = 0.009 and 0.003) and negatively aligned with theta power. In the older group, significant loadings were instead distributed across all five cell stains, two of three rsfMRI metrics (IntTS and ReHo, most strongly ReHo) and low-frequency power (delta and theta). The spatial association between CBF and Parvalbumin staining was retained (r = 0.45 young, 0.36 older; spin-null FDR = 0.040 and 0.047), as was the negative CBF-theta relationship, whereas the spatial associations between CBF and beta- and gamma-band power were no longer detectable (r = 0.34 and 0.31; spin-null FDR = 0.232 for both). The vertex-wise correlations in **Fig. 3d,e** was broadly consistent with the LC1 loadings, with the notable exception of ReHo. ReHo correlated positively with mvPI in both groups (r = 0.75 young, 0.82 older; spin-null FDR < 0.001) and this correlation did not differ between groups (Δr = 0.07, permutation p = 0.136), indicating that its increased LC1 loading in the older group reflects the reweighting of LC1 towards the mvPI pole rather than a change in the underlying vascular-functional relationship. LC1 loadings should therefore be interpreted alongside the bivariate correlations rather than in isolation. Parvalbumin-positive interneurons and high-frequency oscillatory activity are both metabolically demanding^55,56^, and their distributions are elevated in the posterior hippocampus, where we observed the steepest age-related CBF reduction (**Fig. 2e**). In young adults, the spatial distribution of CBF thus tracked markers of high local energy demand, whereas in older adults it aligned more closely with cytoarchitectonic maps. Thus, the spatial correspondence between CBF and high-frequency activity was detectable in young but not older adults.

**Fig. 3.**
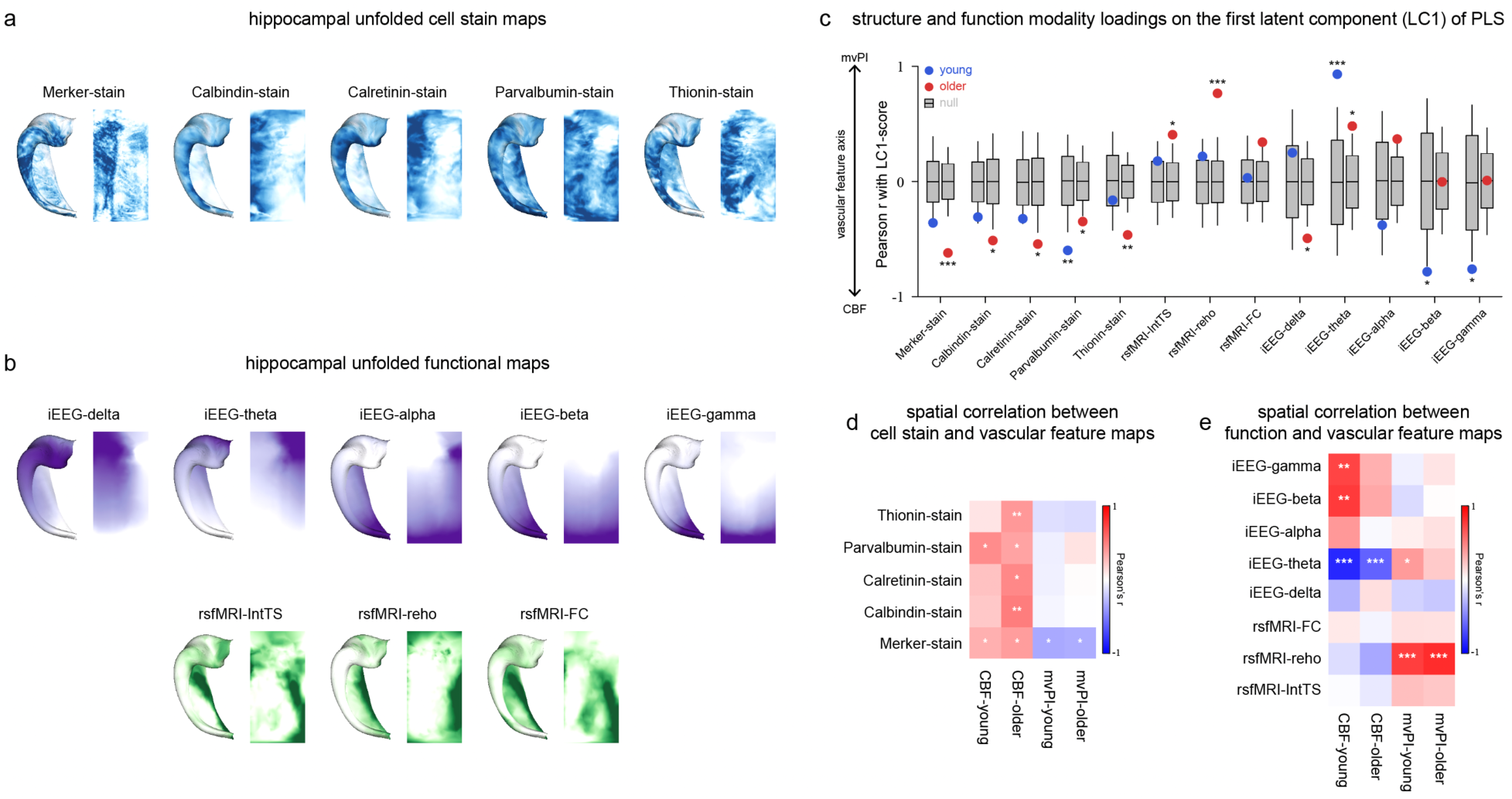
Perfusion and pulsatility show distinct spatial relationships with hippocampal neural organization. **a**, Cell-stain reference maps from HippoMaps (Merker, Calbindin, Calretinin, Parvalbumin, Thionin), each shown on the folded hippocampal surface (left) and in unfolded space (right). **b**, Functional reference maps: band-limited intracranial EEG power (delta, theta, alpha, beta, gamma; purple) and resting-state fMRI measures (intrinsic timescale, regional homogeneity, functional connectivity with cortex; green), shown in the same folded and unfolded views. **c**, Loadings of each reference map on the first latent component (LC1) of a PLS regression relating the two vascular feature maps (CBF, mvPI) to the 13 structural and functional maps. LC1 is oriented such that positive scores correspond to the mvPI pole and negative scores to the CBF pole of the vascular feature axis (left annotation). Grey boxes summarize the null distribution obtained from 10000 spatial surrogate maps generated with spin, which preserve the spatial autocorrelation of each reference map. The horizontal line marks the null mean, box limits mark ± 1 SD, and whiskers span the 2.5th–97.5th percentiles of the null distribution. **d-e**, Vertex-wise Pearson correlations between each reference map and each vascular feature map, shown separately for cell-stain maps (**d**) and functional maps (**e**). Columns give CBF and mvPI in each age group. Asterisks in **c-e** denote significance against the spatial-autocorrelation-preserving (spin) null distribution: *p < 0.05, **p < 0.01, ***p < 0.001, BH-FDR corrected.

### Perfusion and Pulsatility Show Differential Frequency-Specific Relationships with Hippocampal Electrophysiology

The preceding analysis revealed distinct spatial associations of CBF and mvPI with hippocampal electrophysiological organization, particularly the correspondence of mvPI with theta-band power in young adults. Given previous evidence that arterial pulsations can modulate neuronal activity in mice^14^ and that heartbeat-related neural responses in humans often involve theta-range activity^15,21^, we further examined the frequency dependence of these spatial relationships. Specifically, we assessed whether the associations of CBF and mvPI with hippocampal iEEG power were confined to particular frequency bands or extended across a broader spectrum.

We computed spatial correlations between frequency-resolved iEEG power maps (0.5-80 Hz, 0.5 Hz steps) and the CBF and mvPI maps (**Fig. 4a,b**). CBF correlated negatively with iEEG power in the low-to-mid frequency range encompassing the theta band in both age groups. A positive correlation between CBF and high-frequency power was additionally observed in the young group but did not reach significance in the older group. For mvPI, a positive correlation with around theta-range power reached significance only in the young group (**Fig. 4b**). We next compared iEEG power between vertices in the top and bottom deciles of each vascular map (**Fig. 4c,d**, spatial distribution in **Fig. S4**). In the young group, CBF showed greater power in its top decile in the high-frequency range, whereas mvPI showed greater power in its top decile in the theta range. Because CBF and mvPI are spatially anticorrelated (r = -0.37, spin-null p = 0.008, **Fig. 2d**), their decile sets are largely complementary, and these two contrasts are not independent. Neither contrast reached significance in the older group. We next examined whether the observed theta-range pattern could be explained by cardiac contamination of the iEEG signal, such as pulsation-induced electrode motion. Convolving PPG- and ECG-derived kernels with detected R-peaks and computing the resulting power spectrum (**Fig. 4e**) produced a discrete harmonic comb spaced at the mean heart rate (∼1.2 Hz), rather than the broadband elevation across the theta range observed in our data. Thus, the observed spectral pattern differed from that predicted by this simple cardiac-contamination model, for example through pulsation-induced electrode motion. Pulsation-sensitive neurons have been reported in the mouse hippocampus^14^, and heartbeat-locked fluctuations are detectable in human hippocampal iEEG^16,17^. Heartbeat-related MEG responses are concentrated in the theta band^21^, and an intracranial study reported heartbeat-induced phase resetting, rather than amplitude enhancement, of insular theta activity^15^. Cardiac pulsation produces recurring vascular volume changes that may act as rhythmic mechanical inputs to nearby neurons. Such inputs could either evoke additional neural activity or alter the timing of ongoing oscillations without necessarily increasing their amplitude. We therefore simulated three possible ways in which heartbeats could affect neural activity (**Fig. 4f**, details in Methods): each heartbeat could evoke a brief additional response (model A), trigger a short burst of theta-range oscillations (model B), or reset the phase of ongoing theta oscillations without introducing additional oscillatory activity (model C). The three models produced distinguishable spectral signatures (**Fig. 4g**). Model A generated a broad low-frequency elevation; Model B a theta-range increase returning monotonically to baseline; and Model C a sharp theta peak followed by a negative deflection and recovery, reflecting the redistribution of existing power under phase alignment without a net change in total power. This peak-then-undershoot morphology was present in the observed young-group mvPI contrast (**Fig. 4d**) and was qualitatively more similar to the phase-resetting model than to the two additive models. This similarity raises phase resetting as one possible explanation for the observed spectral pattern, but does not establish the underlying mechanism, which will require direct analysis of heartbeat-locked neural phase dynamics.

**Fig. 4.**
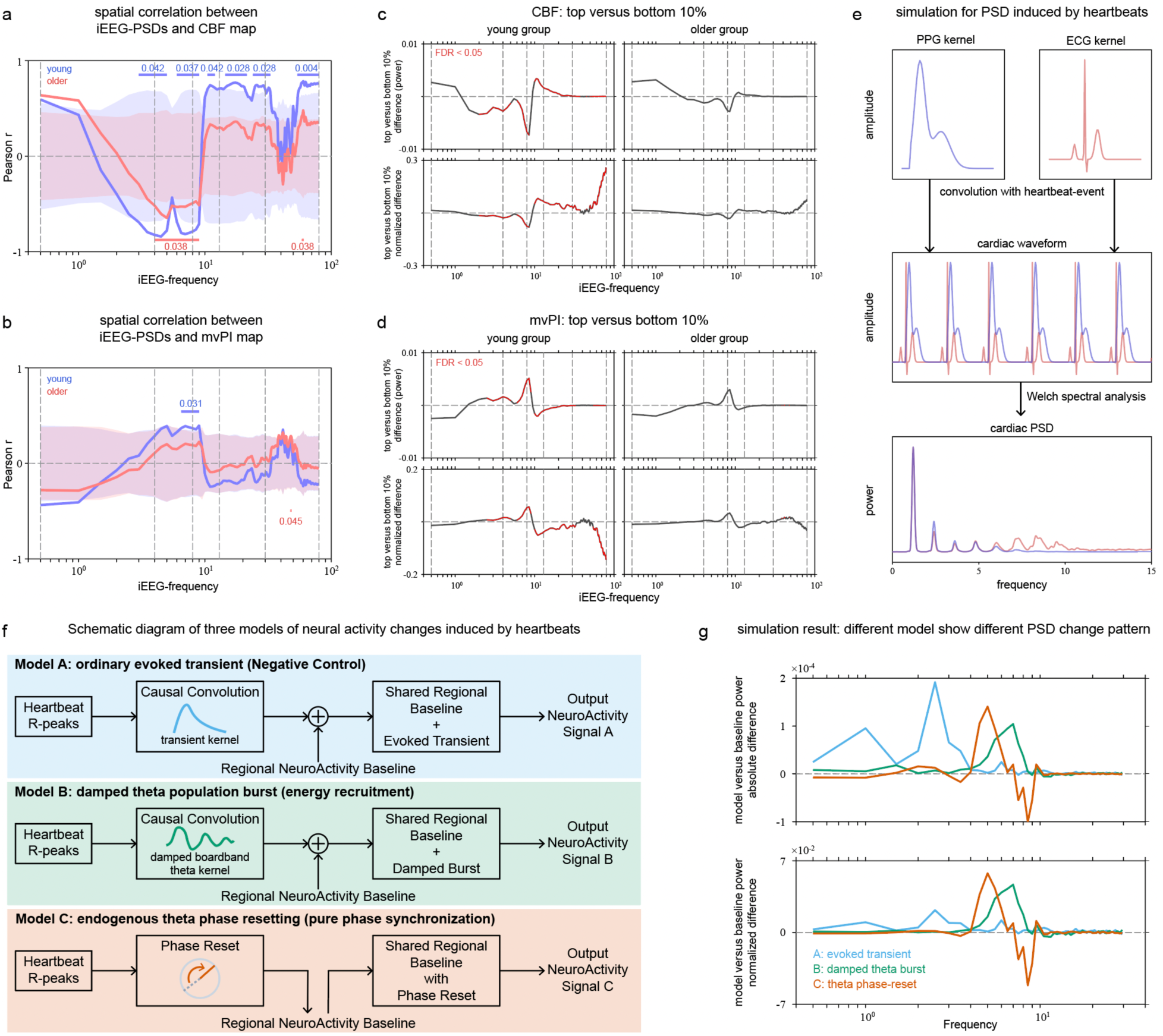
Hippocampal perfusion and pulsatility show distinct frequency-specific relationships with neural activity. **a-b**, Vertex-wise spatial correlation between frequency-resolved iEEG power maps (0.5– 80 Hz, 0.5 Hz steps) and the CBF map (**a**) or the mvPI map (**b**), shown separately for young (blue) and older (red) adults. Shaded bands indicate the 0.5th-99.5th percentiles of null distributions generated from 10000 spatial surrogate maps preserving the spatial autocorrelation of each frequency map. Horizontal bars mark contiguous frequency ranges surviving by permutation cluster test correction across frequency bins, and numbers give the corresponding p values (BH-FDR corrected). **c-d**, Difference in iEEG power spectral density between vertices in the top and bottom deciles of the CBF map (**c**) and the mvPI map (**d**), in young (left) and older (right) adults. Upper rows show absolute power differences, and lower rows show frequency-wise normalized differences, calculated as the difference between the two spectra divided by their mean at each frequency, to emphasize relative changes in spectral shape. Red segments denote frequencies with p < 0.05 (BH-FDR corrected). **e**, Simulation of the spectral signature expected from cardiac contamination of the iEEG signal. PPG- and ECG-derived kernels (top) were convolved with detected R-peak times to synthesis cardiac waveforms (middle), and power spectra were estimated by Welch’s method (bottom). **f**, Three candidate mechanism models by which heartbeats could alter theta-band power, each applied to a shared regional activity baseline. Model A: R-peaks drive a causal convolution with a transient kernel, adding an ordinary evoked response. Model B: R-peaks drive a damped broadband theta burst, representing additive energy recruitment. Model C: R-peaks reset the phase of ongoing endogenous theta without adding power. **g**, Simulated change in power spectral density relative to baseline for each model, expressed as absolute differences (top) and frequency-wise normalized differences (bottom). Model A produces a broad low-frequency elevation, Model B a theta-range increase returning monotonically to baseline, and Model C a sharp theta peak followed by a negative deflection and recovery, the signature of power redistribution under phase alignment with no net change in total power.

Together, these findings reveal distinct spatial relationships between cerebrovascular properties and hippocampal structural and functional organization. In young adults, CBF preferentially corresponded to high-frequency neuronal activity associated with greater metabolic demand, whereas mvPI showed a distinct spatial association with theta-band organization. The frequency-specific mvPI–theta spatial correspondence, together with its qualitative similarity to the phase-resetting simulation, raises heartbeat-related phase resetting as one possible explanation for the observed spectral pattern.

### Spatial Organization of Mechanosensitive Channel Expression along the Perivascular Mechanical Gradient

A previous study has shown that specific pyramidal neurons in the mouse neocortex and hippocampus express stretch-activated, mechanically gated (MG) cation channels that can be activated by membrane suction during cell-attached patch-clamp recordings^12^. Subsequent immunohistochemical study identified Piezo2 expression in select neuronal populations, including neocortical and hippocampal pyramidal neurons^13^. More recently, Salameh et al.^14^ demonstrated that Piezo2 mediates neuronal responses to arterial pulsation in the mouse brain, including the olfactory bulb and hippocampus. Together, these findings identify *PIEZO2* as a candidate molecular substrate through which vascular pulsation could influence neuronal activity. We therefore examined *PIEZO2* expression using the human anterior hippocampal spatial transcriptomic dataset reported by Thompson et al.^51^. We hypothesized that, if *PIEZO2* contributes to the transduction of arterial pulsatility into neuronal activity, its expression might be expected to be spatially associated with the vasculature, with higher expression near blood vessels and a progressive decline with increasing vascular distance. To test this hypothesis, spatial transcriptomic spots were aggregated into distance-stratified pseudo-bulk samples according to their proximity to blood vessels, and *PIEZO2* expression was evaluated across the resulting vascular-distance gradient (G_dist_).

We first performed self-supervised clustering of all spatial transcriptomic spots and annotated the resulting clusters using established marker genes. Vascular masks were then manually delineated on the corresponding histological images combined with annotation, and the distance from each spot to the nearest vessel was calculated (see Methods). **Fig. 5a** and **Fig. S5** show histological images of all 34 captures and the resulting G_dist_, in which spots were classified as vascular (VAS) or into six progressively more distant levels (lvl1-6). Cell-type proportions were estimated using robust cell type decomposition^51,57^ (RCTD), and spots were aggregated into pseudo-bulk samples according to G_dist_. **Fig. 5b** and **Fig. S6** show the distributions of eight major cell types across the vascular-distance gradient. As expected, the vascular-cell proportion was highest in VAS and progressively decreased with increasing G_dist_, supporting the ability of G_dist_ to capture spatial proximity to the vasculature. Several other cell types also exhibited systematic changes across G_dist_. We next examined whether G_dist_ was associated with molecular features related to vascular mechanical properties. After adjustment for estimated vascular-cell proportions, we calculated expression scores for three vascular functional gene sets representing vascular smooth muscle cell (VSMC) contractility, passive mechanical compliance and elastic recoil, and vascular wall stiffness and collagen cross-linking (**Fig. 5c**; see Methods). VSMC contractility and compliance scores decreased significantly from levels 1 to 5, whereas the stiffness score showed a small but significant increase. Accordingly, the balance between compliance and stiffness also decreased significantly with increasing G_dist_ (**Fig. 5d**). Together, these gradients indicate a systematic spatial organization of vascular mechanical-property-related molecular signatures with distance from the vasculature, consistent with spatial variation in the local perivascular mechanical environment. We next asked whether the vascular-distance organization observed in the transcriptomic data could be related anatomically to the spatial organization of hippocampal pulsatility measured with MRI. The schematic in **Fig. 5e** illustrates the predominant arterial supply of the hippocampus^23,24^. Consistent with this anatomy, mvPI showed greater pulsation-related volumetric changes toward the artery-rich medial hippocampal region. Thus, although measured at different spatial scales, the transcriptomic and MRI data both reveal vascular-centered spatial organization: mvPI varies with the distribution of hippocampal arteries at the tissue level, whereas molecular signatures vary with proximity to the vasculature at the microscopic level. Previous work identified Piezo2 as a mediator of neuronal responses to arterial pulsation in mice^14^. Given the slower human heart rate and the broader repertoire of mechanosensitive channels potentially capable of responding to vascular mechanical forces, we extended the analysis to major mechanosensitive channel genes from the PIEZO and K2P families and the TRP superfamily. Because cell-type composition varied systematically across G_dist_ (**Fig. 5b** and **Fig. S6**), we used linear mixed-effects models (LMMs) to assess the association between gene expression and G_dist_ while accounting for the estimated proportions of major cell types, as well as age, sex, capture, and donor effects (see Methods). **Fig. 5f** summarizes the t-statistics for the G_dist_ effect across the examined mechanosensitive channels. *PIEZO2* and *PKD2* showed significant decreases in expression with increasing G_dist_, indicating vascular-distance-associated expression gradients that persisted after accounting for spatial variation in cell-type composition and other measured covariates. Arterial pulsation has also been implicated in driving perivascular cerebrospinal fluid transport within the glymphatic system^9–11^. Consistent with a vascular-associated spatial pattern, *AQP4* expression decreased with increasing G_dist_ (**Fig. 5g**). To determine whether these observations were unusual relative to the transcriptome-wide background, we applied the same G_dist_ analysis to all sufficiently expressed genes and ranked them according to the strength of their distance-associated effects (**Fig. 5h**). *PIEZO2*, *PKD2*, and *AQP4* showed relatively extreme distance-associated effects compared with the transcriptome-wide distribution. Together, these findings demonstrate a vascular-distance-associated spatial organization of *PIEZO2* expression in the human hippocampus, with higher expression near the vasculature and a progressive decline with increasing vascular distance. This spatial pattern is consistent with, but does not establish, a potential role for *PIEZO2* in local vascular mechanotransduction. *PKD2* exhibited a similar spatial organization, identifying it as an additional candidate for vascular-associated mechanosensitive signaling. Exploratory age-stratified analyses further suggested differences in the local perivascular organization of *PIEZO2*, *AQP4*, and vascular mechanical signatures between younger and older samples (**Fig. S7** and **Fig. S8**). However, given the small number of donors and the absence of a significant age-by-distance interaction for *PIEZO2*, these patterns should be considered hypothesis-generating.

**Fig. 5.**
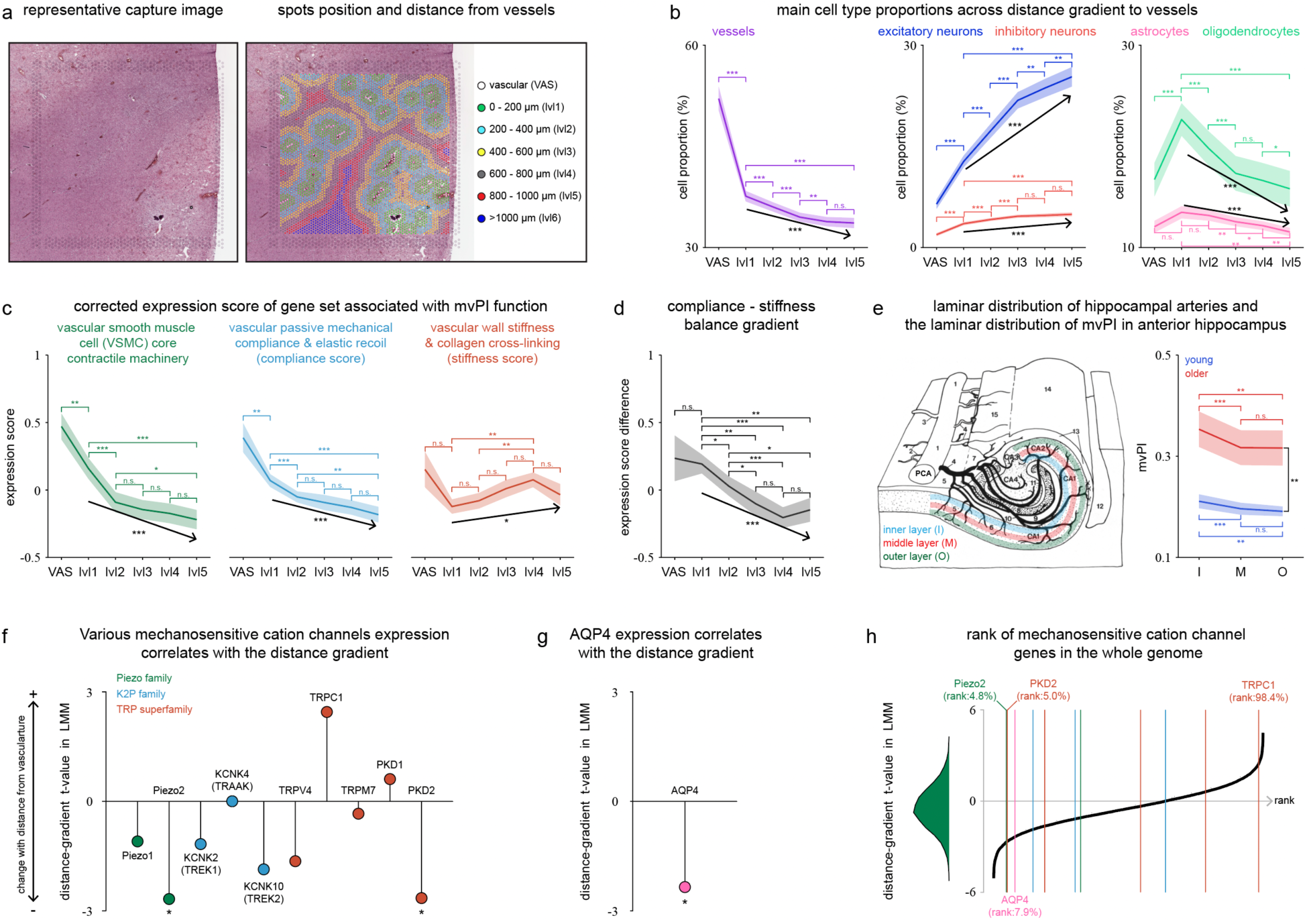
Vascular mechanical signatures and mechanosensitive channel expression are organized along a vascular-distance gradient in the human hippocampus. **a**, Representative capture. Left, Histological staining image. Right, spot map colored by distance-to-vessel bin: vascular ROIs (VAS) manually delineated on the histological image, and concentric 200-μm bins from lvl1 (0–200 μm) to lvl5 (800–1000 μm), with lvl6 (>1000 μm) shown for completeness. **b**, Cell-type proportions across the distance gradient, estimated by RCTD using the matched human hippocampal snRNA-seq dataset from Thompson et al.^51^ as the reference, for vasculature, excitatory and inhibitory neurons, astrocytes and oligodendrocytes (color-coded as in the panel titles). Lines and shaded bands give the mean and s.e.m. across captures. Horizontal bars denote pairwise comparisons between bins (paired T-test, BH-FDR corrected); arrows indicate the monotonic trend across lvl1–lvl5 (regression analysis). *: p < 0.05, **: p < 0.01, ***: p < 0.001, n.s.: not significant, BH-FDR corrected. **c**, Corrected expression scores along the same gradient for three gene sets related to mvPI-relevant vascular function: VSMC core contractile machinery, passive mechanical compliance and elastic recoil (compliance score), and wall stiffness and collagen cross-linking (stiffness score). Scores are corrected for vascular cell proportion. **d**, Difference between the compliance and stiffness scores (compliance − stiffness) across the gradient, indexing the balance between the two. **e**, Laminar organization. Left, schematic of the anterior hippocampus showing the arterial supply and the definition of inner (I), middle (M) and outer (O) layers, adapted by Marinković et al^24^. Right, mvPI by layer in young (blue, n = 15) and older (red, n = 17) adults. Colored bars denote between-layer comparisons within age group, and black bars denote between-age comparisons within layer (unpaired T-test, BH-FDR corrected). **f**, Distance-gradient t values from linear mixed models for mechanosensitive cation channels, grouped by family (Piezo, green; K2P, blue; TRP superfamily, red). Positive values indicate expression increasing with distance from the vasculature. Models included proportions of various cell types within the sample, as well as covariates such as age, sex, capture, and donor. Asterisks denote p < 0.05 after BH-FDR corrected. **g**, As in **f**, for *AQP4*. **h**, Genome-wide calibration of the effects in (**f**) and (**g**). Left, distribution of distance-gradient t values across all 13411 expressed genes; right, the corresponding rank curve, with vertical lines marking the channel genes in f and *AQP4*. Percentile ranks are given for the genes falling in the tails of the genome-wide distribution.

Together, these findings reveal a spatially organized perivascular molecular environment in the human hippocampus, in which vascular mechanical signatures and candidate mechanosensitive pathways vary systematically with vascular proximity. The enrichment of *PIEZO2* near the vasculature further identifies a candidate molecular substrate for local vascular mechanotransduction.

## Discussion

Here, we combined 7T perfusion and microvascular volumetric pulsatility imaging with surface-based hippocampal analysis to characterize two complementary properties of the human hippocampal vasculature: tissue perfusion, indexed by CBF, and pulsation-related vascular volume change, indexed by mvPI. CBF and mvPI showed distinct spatial distributions and age-related patterns across the hippocampus. CBF was lower in older adults across subfields, with the strongest spatially localized difference in the posterior hippocampus, whereas age-related increases in mvPI were spatially heterogeneous. These vascular measures also showed different spatial relationships with hippocampal neural organization. In young adults, CBF preferentially corresponded to parvalbumin-positive interneuron density and beta- and gamma-band activity, whereas mvPI was associated with theta-band organization. Several of these relationships were weaker or no longer detectable in older adults. Complementary analyses provided potential biological context for the pulsatility-associated pattern: its spectral profile was qualitatively similar to that generated by a heartbeat-related phase-resetting model, and spatial transcriptomics revealed vascular-distance gradients of PIEZO2 and other candidate mechanosensitive genes. Together, these findings establish perfusion and tissue-level pulsatility as distinguishable features of the human hippocampal vascular environment and identify distinct spatial relationships with neural organization across age.

The distinct spatial distributions of mvPI and CBF may partly reflect the topology of the hippocampal arterial supply. Anatomical^23,24^ and high-resolution 7T angiographic^22^ studies show that the anterior hippocampal artery (AHA), alone or together with the uncal branch of the anterior choroidal artery (AChA), converges within the uncal and interdigitation sulci of the expanded hippocampal head, forming an extensive superficial anastomotic network. This distribution aligns with the large area of high mvPI and arterial PDD observed in the anterior hippocampus in our data. The relatively long and tortuous course of the arterial branches supplying the anterior hippocampus may increase upstream vascular resistance and reduce the pressure available to the downstream microcirculation, potentially contributing to the relatively low CBF in this region. The middle hippocampal artery (MHA) is constrained by the geometry of the hippocampal sulcus and runs linearly along the longitudinal axis, giving off vertical perforators at right angles, a topology that matches the narrow band of elevated mvPI and arterial PDD along the mid-Sub. The posterior hippocampal artery (PHA) instead enters at multiple points along the caudal curvature from the PCA, the posterior temporal artery and the splenial artery, forming a discrete, comb-like pattern. In contrast, the middle and posterior hippocampus receive arterial branches arising directly from, or relatively close to, the PCA and its major branches, providing a more direct vascular route to the hippocampal body and tail. Together with the denser microvascular networks in CA2/3^25,58,59^, this vascular architecture may contribute to the higher CBF observed in the middle and posterior hippocampus, particularly in CA2/3. Overall, the spatial distributions of CBF and mvPI in the young group are broadly consistent with the topology of the hippocampal arterial supply.

Aging and arteriosclerosis reduce the capacity of central elastic arteries to buffer the cardiac pulse, increasing the pulsatile energy delivered to the cerebral circulation^41^. In the AHA and MHA territories, the age-related elevation of mvPI coincided with the large arteries lying on the hippocampal surface (**Fig. 2c,d**), suggesting that part of the additional pulsatile load may be accommodated by volume excursions of these extraparenchymal segments, thereby attenuating its transmission to the downstream microvasculature. The posterior hippocampus lacks a comparable surface arterial segment, being supplied by smaller, multipoint branches^22^ with low baseline pulsatility and low arterial density (**Fig. 2b,c**), yet showed a pronounced relative increase in mvPI with age (**Fig. 2f**). This pattern raises the possibility that the increased pulsatility occurs predominantly within the parenchymal microvasculature, where less upstream buffering may expose smaller vessels to greater pulsatile load. Sustained microvascular pulsatility has been associated with vascular remodeling, including rarefaction and wall hypertrophy^60–62^, changes that could compromise tissue perfusion. Such processes provide one potential context for the preferential perfusion decline observed in the posterior hippocampus, although their contribution cannot be determined from the present data. Structural and perfusion-related age differences were not spatially coincident throughout the hippocampal long axis: volume reductions were observed in both the middle and posterior hippocampus, whereas significant CBF reduction was concentrated in the posterior hippocampus. The posterior hippocampus therefore showed the clearest spatial overlap between structural and perfusion-related age differences, consistent with previous reports of preferential posterior hippocampal vulnerability during normal aging^52–54^. Regional volume was also positively associated with CBF in the posterior hippocampus. However, given the modest sample size and cross-sectional design, these observations should be considered preliminary and do not establish a temporal or causal relationship between perfusion decline and tissue loss. Larger longitudinal studies will be needed to determine whether these processes are mechanistically related.

The spatial association of CBF with hippocampal cellular and electrophysiological organization is consistent with its role in metabolic substrate delivery. Because the brain has limited endogenous energy reserves, sustained neuronal activity depends on continuous vascular delivery of oxygen and nutrients.

Consistent with this role, CBF preferentially aligned with parvalbumin-positive interneurons and beta- and gamma-band activity in young adults. Fast-spiking PV+ interneurons are particularly metabolically demanding^27,29^ and contribute to fast network oscillations^63^, providing a plausible basis for the correspondence between CBF and these high-energy cellular and electrophysiological features. Notably, this correspondence was selectively altered with age: the CBF–PV relationship was retained, whereas associations with beta- and gamma-band activity were no longer detectable. Together, these findings are consistent with spatial matching between perfusion and the metabolic demands of hippocampal circuits, with the correspondence to fast neuronal activity becoming less apparent with age.

The association of mvPI with theta-band organization raises the possibility that hippocampal vascular pulsatility is related to heartbeat-sensitive neural dynamics. Accumulating evidence supports the capacity of neurons to sense mechanical forces^64,65^. Early studies showed that mechanical stimulation can directly evoke neuronal activity through mechanosensitive ion channels in selected mouse neurons^12^, while subsequent work identified Piezo2 in neuronal populations of the neocortex and hippocampus^13^. More recently, Salameh et al.^14^ demonstrated that arterial pulsations can directly modulate neuronal activity through Piezo2, establishing a mechanistic link between vascular pulsatility and neuronal activity. In humans, heartbeat-related neural responses have been detected with intracranial electrophysiology^15–17^, EEG^19,20^ and MEG^21^, particularly within the theta range, although their contribution from autonomic inputs versus local mechanical transduction remains unresolved. Together, these prior observations provide a mechanistic context for considering whether the mvPI–theta spatial correspondence observed here could be related to local vascular mechanotransduction. Notably, our simulations were more consistent with heartbeat-induced phase resetting^15,66^ than with additive neuronal activation. This similarity raises phase resetting as one possible explanation for the observed spectral pattern, although the simulations do not establish the underlying physiological mechanism. Complementing this observation, spatial transcriptomic analysis revealed a vascular-distance gradient of *PIEZO2* expression in the human hippocampus, with higher expression near the vasculature. Given the established role of Piezo2 in arterial-pulsation-sensitive neurons in mice^14^, this spatial organization identifies *PIEZO2* as a candidate molecular substrate for local vascular mechanotransduction in the human hippocampus. *PKD2* showed a similar spatial gradient, identifying an additional candidate mechanosensitive pathway, potentially related to previously described astrocyte-mediated infra-slow oscillation vascular–neural signaling^67^. Together, these observations raise local vascular mechanotransduction as one candidate explanation for the association between tissue-level pulsatility and theta-band organization observed here, but direct temporal and experimental measurements will be required to determine whether such a pathway operates in humans. Exploratory analyses further suggested age-related differences in perivascular molecular signatures, including the spatial organization of *PIEZO2* and *AQP4*. However, given the limited donor number and the absence of a significant age-by-distance interaction for *PIEZO2*, these findings should be considered hypothesis-generating and require validation in larger datasets.

Several limitations should be considered. First, the present study is cross-sectional and relies largely on spatial correspondence across multimodal datasets. The observed vascular–neural associations therefore cannot establish causal or directional relationships. In particular, direct heartbeat-resolved measurements of vascular and neural dynamics will be required to determine whether local vascular pulsatility is temporally related to neuronal activity and, if so, whether phase resetting contributes to this relationship. The modest imaging sample size, particularly after stratification by age, also limits the precision of estimated age-group differences and warrants replication in larger cohorts. Second, the histological, electrophysiological, and transcriptomic reference datasets were acquired from cohorts independent of our MRI participants. The iEEG maps were derived primarily from younger patients with epilepsy and may therefore not fully represent the electrophysiological organization of the healthy aging hippocampus. Consequently, these analyses establish group-level spatial correspondence rather than within-subject vascular–neural relationships; simultaneous or within-subject vascular and electrophysiological measurements will be needed to test whether these relationships are also expressed at the individual level. Finally, the spatial transcriptomic analysis was limited by the small number of donors and incomplete characterization of the local vascular mechanical environment. Vascular distance was estimated within two-dimensional sections, omitting out-of-plane vessels, and distance itself is only an indirect proxy for mechanical exposure because pulsatility varies across vessels and anatomical locations. These factors may increase heterogeneity across captures and limit attribution of the observed molecular gradients specifically to pulsatile forces. The future studies combining three-dimensional vascular reconstruction with local measurements of pulsatility will be needed to characterize the perivascular mechanical–molecular interface more directly.

In conclusion, our findings identify perfusion and tissue-level vascular pulsatility as distinct features of the human hippocampal vascular environment, with different spatial organization, age-related patterns, and relationships with neural architecture. Perfusion preferentially corresponded to cellular and electrophysiological features associated with high metabolic demand, whereas pulsatility showed a distinct spatial relationship with theta-band organization. The latter, together with vascular-distance gradients of candidate mechanosensitive genes, raises local vascular mechanotransduction as a hypothesis for future direct testing. By integrating vascular imaging with electrophysiological and molecular spatial maps within a common anatomical framework, this study provides a general strategy for investigating vascular–neural relationships beyond the hippocampus. More broadly, our findings position tissue-level vascular pulsatility alongside perfusion as a potentially informative dimension of the human cerebrovascular environment whose relationship with neural organization warrants investigation across the brain.

## Methods

### Ethics statement

This study complied with all relevant ethical regulations. The study protocol was approved by the Institutional Review Board (IRB) of the University of Southern California. All participants provided written informed consent prior to participation. Participant sex was determined based on self-report. The study included 20 female and 15 male participants. Sex was not considered as an experimental factor.

### Participants

15 young (mean age = 29.7 years, SD = 4.8 years, 6 females) and 20 older (mean age = 71.1 years, SD = 9.3 years, 14 females) volunteers participated in this study. All participants in this study were healthy without known neurological, psychiatric disorders or major systemic diseases and refrained from caffeine 3 hours before the scan. All participants provided written informed consent according to a protocol approved by the IRB of the University of Southern California and were paid for attendance (40 US dollars). Head motion was minimized by placing cushions on top and two sides of head. Due to technical or personal reasons, some older participants did not complete the full set of scans. Specifically, 18 completed ASL, 18 completed VASO, and 16 completed TOF. In the final analysis, data were retained based on a combination of head motion and HippUnfold segmentation results: 17 ASL, 17 VASO, and 16 TOF.

### MRI data acquisition

MRI data were acquired on the investigational pTx part of a 7T Terra system (Siemens Medical Systems, Erlangen, Germany) with an 8Tx/32Rx head coil (Nova Medical, Cambridge, MA, USA). Anatomical images were acquired with a T1-weighted MP2RAGE sequence (0.7-mm isotropic voxels, 224 sagittal slices, FOV = 224×224 mm, TR = 4500 ms, TE = 3.43 ms, TI1 = 1000 ms, flip angle = 4°, TI2 = 3200 ms, flip angle = 5°, bandwidth = 200 Hz/Px, slice partial Fourier = 6/8, GRAPPA = 3).

Functional images were acquired for resting states with a slice-selective slab-inversion (SS-SI) vascular space occupancy (VASO) sequence^48–50^ with following imaging parameters: 1.2 mm isotropic voxels, 28 axial slices, TI1/TI2/shotTR/TE = 1,355/2,475/40.0/10.3 ms, partial Fourier factor = 6/8, variable flip angle scheme with reference flip angle = 33°, three-fold GRAPPA acceleration in z direction with a CAIPIRINHA shift of 1, bandwidth = 1,420 Hz/Px, FOV = 153.6 × 192 mm, matrix size = 128 × 160 and acquisition time approximately 20 minutes with concurrent pulse recording.

For the time-of-flight angiography (TOF-MRA), with following imaging parameters: 0.5 mm isotropic voxels, 3 slab with 40 slices, TR/TE = 19.0/3.61 ms, flip angle = 24°, GRAPPA = 3, bandwidth = 187 Hz/Px, FOV = 200 × 200 mm, matrix size = 400 × 400 and acquisition time 6:26.

For the perfusion image, a whole-brain 3D pCASL imaging with 1 mm isotropic resolution was performed at 7T using the FLASH-based rotated golden-angle stack of spirals (rGA-SoS)^46,47^ sampling scheme with following imaging parameters: FOV = 256 × 256 × 144 mm^3^, matrix size = 256 × 256 × 144, TR = 7 s, echo spacing (ES) = 19.5 ms, TE = 1.72 ms, ADC bandwidth = 400k Hz, ADC duration = 10.5 ms, flip angle = 15°, and echo train length (ETL) = 72, total acceleration factor (R) = 4. Each M0, label, and control acquisition was acquired over 12 TRs, which was divided into 3 dynamic frames and can be reconstructed into 3 individual images to avoid lengthy temporal footprint, resulting in an effective temporal resolution for each image of 28 sec. In each TR, 1 spiral interleave with variable density (R_spiral_ = 2) in the kxy plane was acquired across 72 kz samples (Rz = 2). GA rotation (137.5°) was applied to spiral interleaves across 3D spatial space and dynamic frames, enabling superior incoherence in 4D. pCASL labeling has a duration of 1500 ms, and post labeling delay was 1800 ms. To avoid the loss of labeling efficiency due to field inhomogeneity at 7T. An optimized parameter set (Optimization of pseudo-continuous arterial spin labeling at 7T with parallel transmission B1 shimming^68^) and a phase adjustment according to a prescan at the beginning of each scan (optimization of pseudo-continuous arterial spin labeling using off-resonance compensation strategies at 7T) was used.

### ASL reconstruction

We used a novel approach for ASL by integrating compressed sensing (CS) reconstruction with motion-resolved self-navigation to minimize motion artifacts^69^. Dynamic CS reconstruction incorporating spatial regularization on total variation (TV_r_) and temporal regularization on finite difference (FD_t_) across dynamic frames is performed to exploit the sparsity in 4D enabled by the sequence, which is described by,

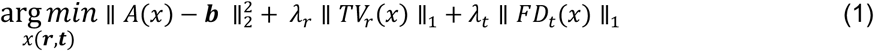

where x and b represent 4D image and k-space data, respectively. The forward operator A incorporates coil sensitivity weighting, non-uniform fast Fourier transform (NUFFT), and k-space undersampling. The inverse problem was solved using alternating direction method of multipliers (ADMM). Reconstruction was performed using 10 ADMM iterations with 3 nested conjugate gradient steps per iteration. Blurring caused by spiral sampling was mitigated by multi-frequency interpolation (MFI), and spiral trajectory errors were corrected by incorporating gradient impulse response (GIRF) that was measured on the scanner.

### MRI data preprocessing, hippocampus segmentation and unfolding

MRI data preprocessing were conducted using AFNI^70^, FreeSurfer^71^, ANTs^72^ and MATLAB (R2021a). The preprocessing of the ASL data is following linear motion correction for control and label images, and registration average control image to T1w anatomical image. The perfusion image obtained by subtracting the label image from the control image can be used to obtain the CBF image according to the classic pCASL calculation method^73,74^. The preprocessing of the VASO data includes removing spikes in time series, linear motion correction and registration average corrected VASO image to T1w anatomical image. Microvascular volumetric pulsatility index (mvPI) was calculated as previously described^48^. Briefly, VASO images were retrospectively sorted into cardiac phases according to the simultaneously recorded pulse signal. Cardiac-cycle CBV changes (ΔCBV) were derived from the VASO signal fluctuations, and baseline CBV (CBV_0_) was estimated from ASL-derived CBF using the Grubb law^75^ (with assuming the average baseline CBV_0_ is 5.5% in cortical gray matter^76^). The mvPI was then calculated following:

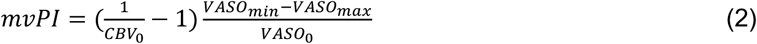

The preprocessing of the TOF-MRA images were registered to the T1w anatomical image, followed by Rician noise denoising, N4 bias-field correction, and vessel enhancement using a multiscale Hessian-based vesselness filter^77^.

The hippocampus was segmented from the high-resolution T1w anatomical images using HippUnfold^36^, which uses hippocampal topology and Laplacian-based unfolding to establish an intrinsic coordinate system for subfield segmentation and surface reconstruction. The preprocessed imaging measures were subsequently mapped onto the hippocampal surface using the HippoMaps framework^37^ for subsequential analysis. Despite limited spatial resolution for resolving individual hippocampal subfields and layers, sampling a large number of voxels across varying depths and subfields provided robust signal estimates, substantially attenuating partial volume artifacts^49,78–81^ (ROI-based analysis).

### Spin-null model

Statistical significance of spatial correlations between hippocampal surface maps was assessed using the spin-test implemented in HippoMaps^37^. To account for spatial autocorrelation, 10,000 spatially constrained permutations were generated by randomly rotating the surface maps while preserving their spatial organization. The observed correlation was compared with the resulting null distribution to obtain a spatial-autocorrelation-corrected p value.

### Hippocampal reference maps

Multimodal hippocampal reference maps were obtained from the HippoMaps repository, in which datasets from different modalities are spatially aligned to a common hippocampal surface coordinate system using HippUnfold. The selected maps comprised five ex vivo histological stains (Merker, Calbindin, Calretinin, Parvalbumin and Thionin), three resting-state fMRI measures (intrinsic timescale, regional homogeneity and hippocampal–neocortical functional connectivity), and intracranial EEG power in five frequency bands (delta, theta, alpha, beta and gamma). Histological maps were derived from three-dimensional ex vivo datasets^82,83^, rsfMRI maps from group-averaged resting-state data in healthy participants^37^, and iEEG maps from hippocampal recordings in patients with epilepsy^37,84^, with sparse electrode measurements spatially mapped to the hippocampal surface according to geodesic distance. All reference maps were represented in the standardized HippoMaps surface space for subsequent spatial analyses.

### Partial least squares regression

Partial least squares (PLS) regression was used to identify latent spatial patterns linking cerebrovascular measures to hippocampal structure–function organization. The predictor matrix X comprised selected hippocampal reference maps from HippoMaps, while the response matrix Y comprised the group-averaged CBF and mvPI maps. All maps were z-score standardized across hippocampal surface vertices, and PLS regression was performed separately for the young and older groups. Conceptually, the cross-block covariance structure can be expressed by singular value decomposition:

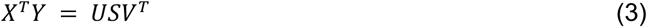

where U and V represent the weights of the hippocampal reference maps and cerebrovascular measures, respectively, and the diagonal elements of S quantify the covariance captured by each latent component. The association of each hippocampal reference map with the first latent component (LC1) was quantified by the Pearson correlation between the reference map and the corresponding LC1 spatial score:

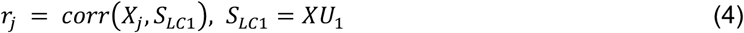

where X_j_ denotes each hippocampal reference map, and S_LC1_ denotes the spatial score of the first latent component. Statistical significance of these correlations was assessed using 10,000 spatial permutations generated by the HippoMaps spin-null model to account for spatial autocorrelation, with multiple comparisons corrected using the Benjamini–Hochberg false discovery rate (BH-FDR).

### Simulation of heartbeat-related neural modulation

To examine whether the observed theta-range spectral pattern could arise from different forms of heartbeat-related neural modulation, we performed forward simulations of three candidate mechanisms. A common 300 seconds synthetic iEEG background was generated from multiple oscillatory components approximating the spectral organization of the hippocampal reference data. Heartbeat events were generated at a mean rate of 72 beats per minute with low-frequency and respiratory modulation of the R–R intervals and additional beat-to-beat variability. The same heartbeat sequence, beat amplitudes, background activity and phase perturbations were used across all models to isolate differences arising from the modulation mechanism.

Three models were constructed to represent different ways in which heartbeats might influence ongoing neural activity (**Fig. 4f**). In Model A (evoked response), each heartbeat generated a brief additional signal, analogous to a transient neural response triggered by an external stimulus. This response was added to the ongoing background activity. In Model B (induced oscillatory burst), each heartbeat triggered a short period of theta-range oscillations that gradually decreased in amplitude over time. Unlike Model A, which introduced a brief transient, Model B added several cycles of oscillatory activity. In Model C (phase resetting), heartbeats did not generate additional signals. Instead, each heartbeat shifted the phases of existing oscillatory components toward preferred phases, temporarily aligning their timing while preserving their amplitudes. Thus, Models A and B simulated heartbeat-related additions to ongoing neural activity, whereas Model C simulated a change in the temporal organization of existing activity without introducing additional oscillatory power. Power spectral density (PSD) was estimated using Welch’s method with an 8-s Hamming window, 50% overlap and 4,096-point FFT, after excluding 5 s from each end of the simulated time series. Spectra were subsequently averaged into 0.5-Hz bins. Model-induced spectral changes were evaluated relative to the common baseline spectrum using both absolute PSD differences and normalized differences.

### Human hippocampal spatial and single-nucleus transcriptomic datasets

Human hippocampal spatially resolved transcriptomic (SRT) and single-nucleus RNA-sequencing (snRNA-seq) data were obtained from the atlas generated by Thompson et al.^51^. The datasets were derived from postmortem anterior hippocampal tissue from 10 neurotypical adult donors, with SRT and snRNA-seq measurements obtained from adjacent tissue sections. SRT data were generated using the 10x Genomics Visium Spatial Gene Expression platform on 10-μm H&E-stained tissue sections. The original study generated 36 Visium-H&E capture areas (2–5 per donor). Of these, data from 34 capture areas that were publicly available to us were included in the present analysis. The spatial coordinates, gene-expression matrices and corresponding histological images provided with the published dataset were used for subsequent spatial analyses. The accompanying snRNA-seq dataset was generated using the 10x Genomics Chromium platform from adjacent hippocampal sections. For each donor, nuclei were collected using PI^+^ sorting and neuronal-enriched PI^+^/NeuN^+^ sorting, yielding 26 snRNA-seq libraries across the 10 donors.

### SRT clustering and annotation

Spatial transcriptomic spots were clustered and annotated following the workflow described by Thompson et al.^51^. Briefly, gene-expression data were normalized and subjected to dimensionality reduction, followed by self-supervised clustering to identify spatially and transcriptionally coherent spot clusters. Cluster identities were assigned based on the expression of established hippocampal cell-type and anatomical marker genes, together with their spatial distribution on the corresponding H&E-stained tissue sections. The resulting annotations were used in combination with histological information to guide the identification of vascular regions.

### Cell-type deconvolution of SRT data

Cell-type composition of the spatial transcriptomic spots was obtained from the RCTD deconvolution^57^ performed by Thompson et al.^51^. Briefly, Thompson et al. evaluated three spatial deconvolution approaches (cell2location, Tangram and RCTD) using their matched hippocampal snRNA-seq dataset as the single-cell reference. Deconvolution performance was evaluated against cell counts derived from immunofluorescence imaging, with RCTD showing the highest concordance with the imaging-based estimates. RCTD was therefore applied to the Visium-H&E dataset using its default marker-gene selection strategy. We directly used the publicly available spot-level RCTD results provided by Thompson et al. through their accompanying GitHub repository. For each spatial transcriptomic spot, the estimated proportions of the annotated cell types were extracted and subsequently used to characterize changes in cellular composition across the vascular-distance gradient and as covariates in analyses of distance-dependent gene expression.

### Vascular-distance mapping and distance-stratified pseudo-bulk analysis

Vascular regions of interest (VAS ROIs) were manually delineated for each spatial transcriptomic capture on the corresponding H&E-stained tissue image, with the spatial domain annotations overlaid as anatomical guidance. Multiple vascular ROIs could be delineated within each capture and were combined into a binary vascular mask. The Euclidean distance from every image pixel to the nearest vascular-mask boundary was calculated and converted from image pixels to physical distance (μm) using the Visium spot diameter (55 μm) and the image-specific spatial scaling factor. The distance assigned to each transcriptomic spot was determined from the distance map at its spot-center coordinate.

Spots located within the manually delineated vascular mask were classified as the vascular region (VAS; distance = 0). Remaining spots were stratified according to their distance from the nearest vascular ROI into consecutive 200-μm bins: lvl1 (0–200 μm), lvl2 (200–400 μm), lvl3 (400–600 μm), lvl4 (600–800 μm), lvl5 (800–1000 μm), and lvl6 (>1000 μm). Together, these categories defined the vascular-distance gradient (G_dist_). Spot coordinates, raw gene-expression counts, and the published RCTD cell-type proportion estimates were matched by spatial barcode. RCTD weights were normalized to sum to one within each spot; cell-type contributions below 1% were set to zero and the remaining weights were renormalized. Spots lacking matched gene-expression or RCTD information were excluded from subsequent analyses.

For distance-stratified pseudo-bulk analysis, spots were aggregated separately within each G_dist_ bin and each capture. Spots with an estimated CSF-associated cell proportion ≥50% were excluded to minimize contamination from choroid plexus or CSF-dominated regions. Within each capture and distance bin, raw expression counts were averaged across the retained spots to generate a capture-level expression profile; RCTD-estimated cell-type proportions were averaged in the same manner. Expression values were converted to counts per million (CPM), with mitochondrial and hemoglobin genes excluded from the library-size denominator and transformed as log_2_(CPM+1) for subsequent gene-expression analyses. Unless otherwise specified, capture-level pseudo-bulk profiles were retained as the primary unit of analysis, preserving capture-specific spatial variation while allowing donor identity to be incorporated in subsequent statistical models.

### Vascular mechanical and mechanosensitive gene panels

Three compact vascular gene panels were defined to represent core molecular features of vascular mechanics: VSMC contractility^85^ (ACTA2, MYH11, TAGLN, CNN1, and MYLK), passive compliance and elastic recoil^86^ (ELN, FBN1, FBLN5, FBLN2, EFEMP1, MFAP5, FBLN1, LTBP4, and DCN), and vascular wall stiffness and ECM cross-linking^87^ (COL1A1, COL1A2, COL3A1, COL5A1, COL5A2, LOX, LOXL1, and LOXL2). These panels were intentionally restricted to core genes with established roles in smooth-muscle contraction, elastic-fiber organization, or collagen-based wall stiffness to provide specific and interpretable molecular proxies of vascular mechanical properties. A mechanosensitive channel panel comprising PIEZO1, PIEZO2, KCNK2 (TREK1), KCNK4 (TRAAK), KCNK10 (TREK2), TRPV4, TRPC1, TRPM7, PKD1, and PKD2 was similarly selected to represent major channel families implicated in membrane tension, stretch, or mechanically coupled signaling^14,88^.

### Vascular mechanical gene-set scores

Gene-set scores for VSMC contractility, passive compliance and elastic recoil, and vascular wall stiffness were calculated from the distance-stratified pseudo-bulk expression profiles. For each gene, expression values across all captures and distance bins were standardized using a global z-score, and the mean z-scored expression of genes within each panel was used as the corresponding raw gene-set score. Genes with low mean expression (mean log_2_CPM < 2) or negligible variance were excluded. To account for differences in vascular-cell abundance across spatial samples, each raw gene-set score was linearly regressed against the estimated vascular-cell proportion. The vascular-adjusted score was obtained by removing the fitted vascular-cell component while preserving the overall score scale through mean-centering of the vascular-cell proportion. Spatial trends across the vascular-distance gradient were assessed across lvl1–lvl5 by fitting a linear regression within each capture and testing the resulting slopes against zero across captures, with pairwise differences between distance bins assessed using paired t-tests and BH-FDR correction.

### Linear mixed-effects modelling of G_dist_-associated gene expression

Linear mixed-effects models (LMMs) were used to assess gene-expression changes along the vascular-distance gradient while accounting for spatial variation in cell-type composition and the nested structure of the dataset. Analyses were restricted to lvl1–lvl5, excluding the manually delineated VAS region, and distance was represented by the centers of the 200-μm bins. Gene expression, distance and RCTD-estimated cell-type proportions were z-score standardized before model fitting. For each gene, the model was specified as:

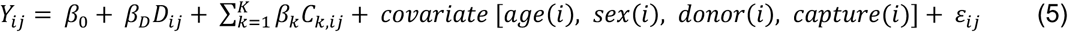

where Y_ij_ denotes standardized gene expression in distance bin j of capture i, D_ij_ denotes vascular distance, and C_k,ij_ denotes the RCTD-estimated proportions of major cell classes. Capture, donor, age and sex were included as covariate terms. OPC proportion was omitted because of its low abundance and to avoid collinearity among compositional covariates, thereby ensuring a full-rank design matrix. The t-statistic and corresponding two-sided p value for β_D_ were used to quantify the direction and strength of the vascular-distance association. For the predefined mechanosensitive channel panel, p values were corrected across genes using the BH-FDR.

### Statistical analysis

Statistical analyses were performed in MATLAB (R2021a). Unless otherwise specified, all statistical tests were two-sided and statistical significance was defined as p < 0.05. Multiple comparisons within each predefined family of tests were controlled using the Benjamini–Hochberg false discovery rate (BH-FDR). Differences in hippocampal CBF and mvPI across age groups and subfields were assessed using 2 (age group) × 7 (subfield) mixed-design repeated-measures ANOVAs, with age group as the between-participant factor and subfield as the within-participant factor. Greenhouse–Geisser correction was applied where appropriate, and effect sizes are reported as partial η^2^. Post hoc comparisons between age groups within individual subfields were performed using two-sample t-tests with BH-FDR correction across subfields. Spatially resolved between-group differences across the unfolded hippocampal surface and along the anterior–posterior axis was assessed using permutation-based tests, with multiple comparisons corrected as specified for each analysis. Spatial correlations between hippocampal maps were quantified using Pearson correlation, and statistical significance was evaluated against 10,000 spatial surrogate maps generated using the HippoMaps spin-null procedure to preserve spatial autocorrelation. Resulting p values were BH-FDR corrected across the corresponding set of comparisons. For spatial transcriptomic analyses, comparisons between vascular-distance bins within the same capture were performed using paired t-tests. Monotonic trends across lvl1–lvl5 were assessed by fitting a linear regression within each capture and testing the resulting slopes against zero across captures. Gene-level associations with vascular distance were assessed using linear mixed-effects models as described above, with significance of the distance effect evaluated from the corresponding model coefficient. For the predefined mechanosensitive channel panel, p values were BH-FDR corrected across tested genes. Age-stratified VAS–lvl1 comparisons were performed using paired t-tests within each age group, whereas between-age comparisons were performed using two-sample t-tests. Age × spatial-region interactions were assessed using mixed-design repeated-measures ANOVA. Given the limited number of donors in the age-stratified transcriptomic analysis, these analyses were considered exploratory. Unless otherwise stated, summary data are presented as mean ± s.e.m.

## Data Availability

The MP2RAGE, ASL, VASO and TOF-MRA datasets generated in this study are made publicly available in the OpenNeuro repository under accession [ds008942]. Source Data underlying the figures are provided with this paper. Multimodal hippocampal reference maps, including histological, resting-state fMRI and intracranial EEG maps, were obtained from the publicly available HippoMaps^37^ resource (https://hippomaps.readthedocs.io/en/latest/). Human hippocampal spatial transcriptomic and single-nucleus RNA-sequencing data were obtained from Thompson et al.^51^ and are publicly available through the Gene Expression Omnibus under accession GSE264692 (SRT) and GSE264624 (snRNA-seq).

## Code Availability

The custom analysis and plotting code developed in this study is publicly available at GitHub (https://github.com/FanhuaGuo/VascularMRI-Framework).

## Acknowledgements

The authors sincerely thank all study participants for their time and contribution to this research. We also thank the developers and contributors of HippUnfold and HippoMaps and the human hippocampal spatial transcriptomic resources provided by the spatial_hpc project for making these valuable datasets, analytical tools, and resources publicly available.

## Funding

This work was supported by US National Institute of Health (NIH) grant UF1-NS100614, S10-OD025312, R01-EB032169, RF1-AG084072, R01-MH134004, R01-AG082073, and R01-NS134712.

## Author Contributions

F.G. and D.J.W. conceived the study. F.G. and D.J.W. designed experiments. F.G., A.J.K., C.Z. and Z.L. executed experiments. F.G. provided support to VASO and TOF-MRA. C.Z. provided support to ASL. F.G. and C.Z. contributed to imaging protocol optimization. F.G. analyzed experiments. F.G. and D.J.W. drafted the manuscript. F.G., A.J.K., C.Z., T.D., Z.L., Z.Y., K.J., X.S., N.J., M.M. and D.J.W. interpreted the results, reviewed the paper, and approved the decision to submit the paper.

## Competing Interests

All authors declare no competing interests.

## Supplement Information

**Fig. S1.**
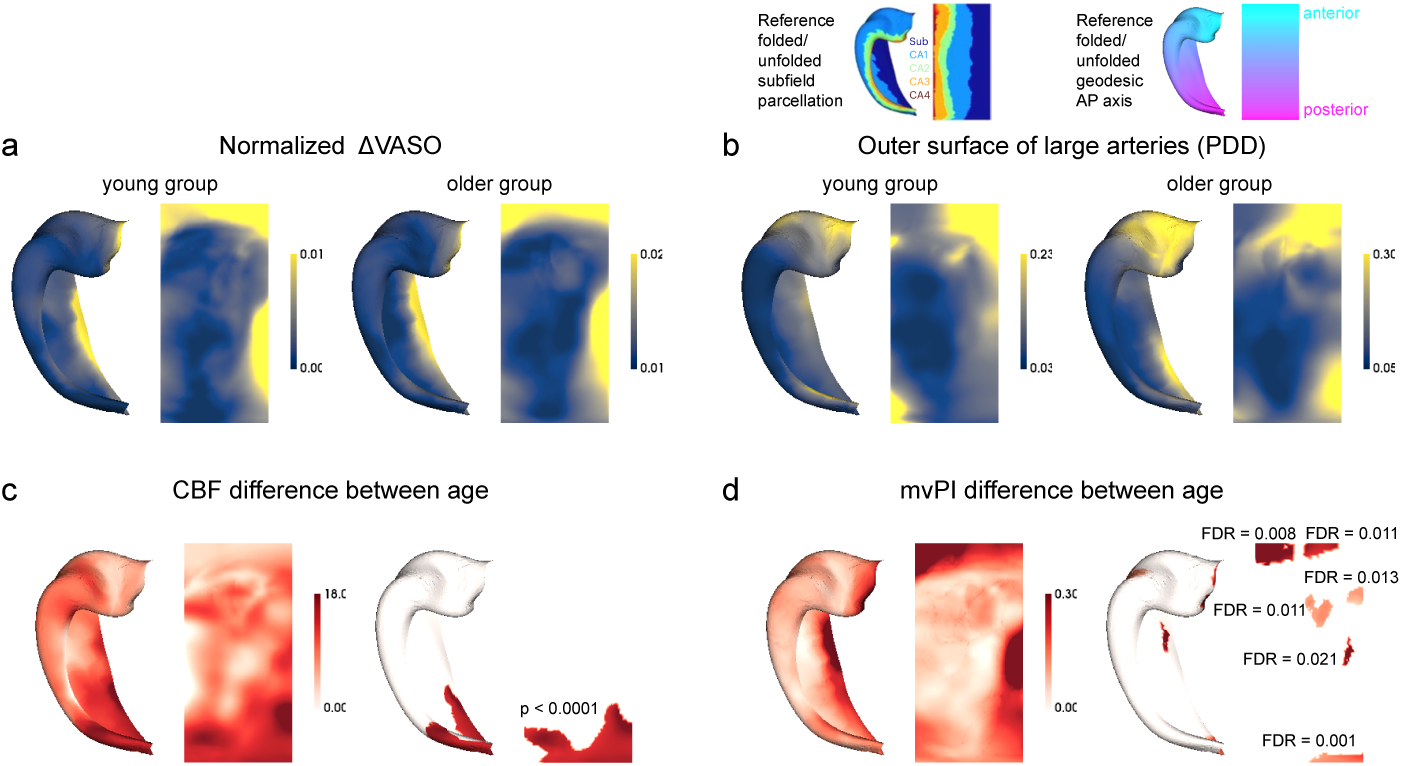
Additional MRI unfolded hippocampal spatial maps. **a**, Group-average normalized ΔVASO maps in unfolded hippocampal space for young (left, n = 15) and older (right, n = 17) adults. **b**, Probability density distribution (PDD) of large arteries (diameter ≥ 0.5 mm) adjacent to the outer hippocampal surface, derived from TOF angiograms and averaged across young (left, n = 15) and older (right, n = 16) adults. Values give the across-participant probability of a large artery being present within each vertex. **c**, age difference in CBF computed vertex-wise. Right panel shows the significant clusters of group differences (permutation cluster test, vertex-wise p < 0.001, BH-FDR correction). **d**, As in **c** for mvPI. structure and function modality loadings on the first latent component (LC1) of PLS

**Fig. S2.**
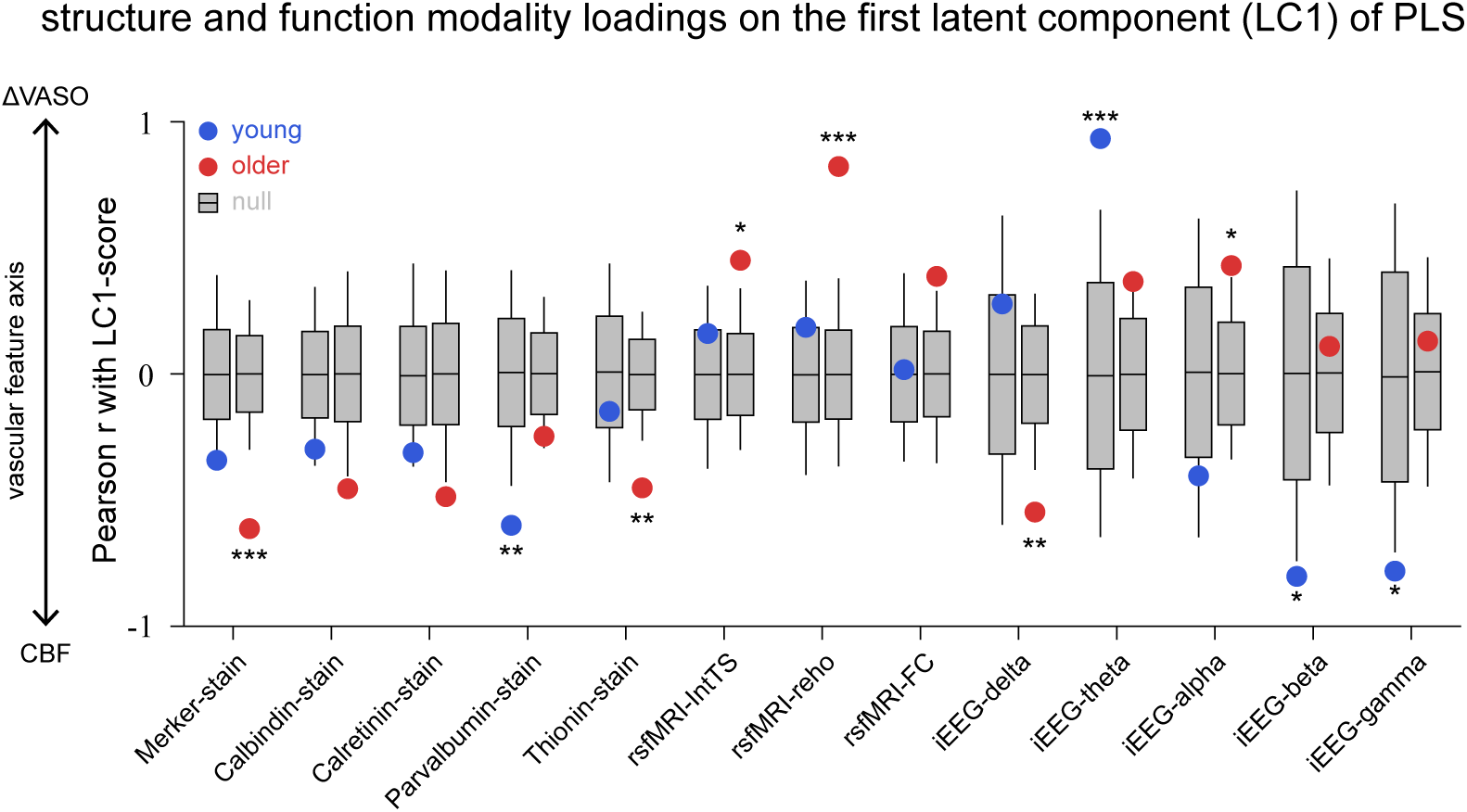
Replicating PLS results using ΔVASO instead of mvPI. Loadings of each reference map on the first latent component (LC1) of a PLS regression relating CBF and ΔVASO to the 13 structural and functional maps. Grey boxes summaries the null distribution obtained from 10000 spatial surrogate maps generated with spin, which preserve the spatial autocorrelation of each reference map. The horizontal line marks the null mean, box limits mark ± 1 SD, and whiskers span the 2.5th–97.5th percentiles of the null distribution.

**Fig. S3.**
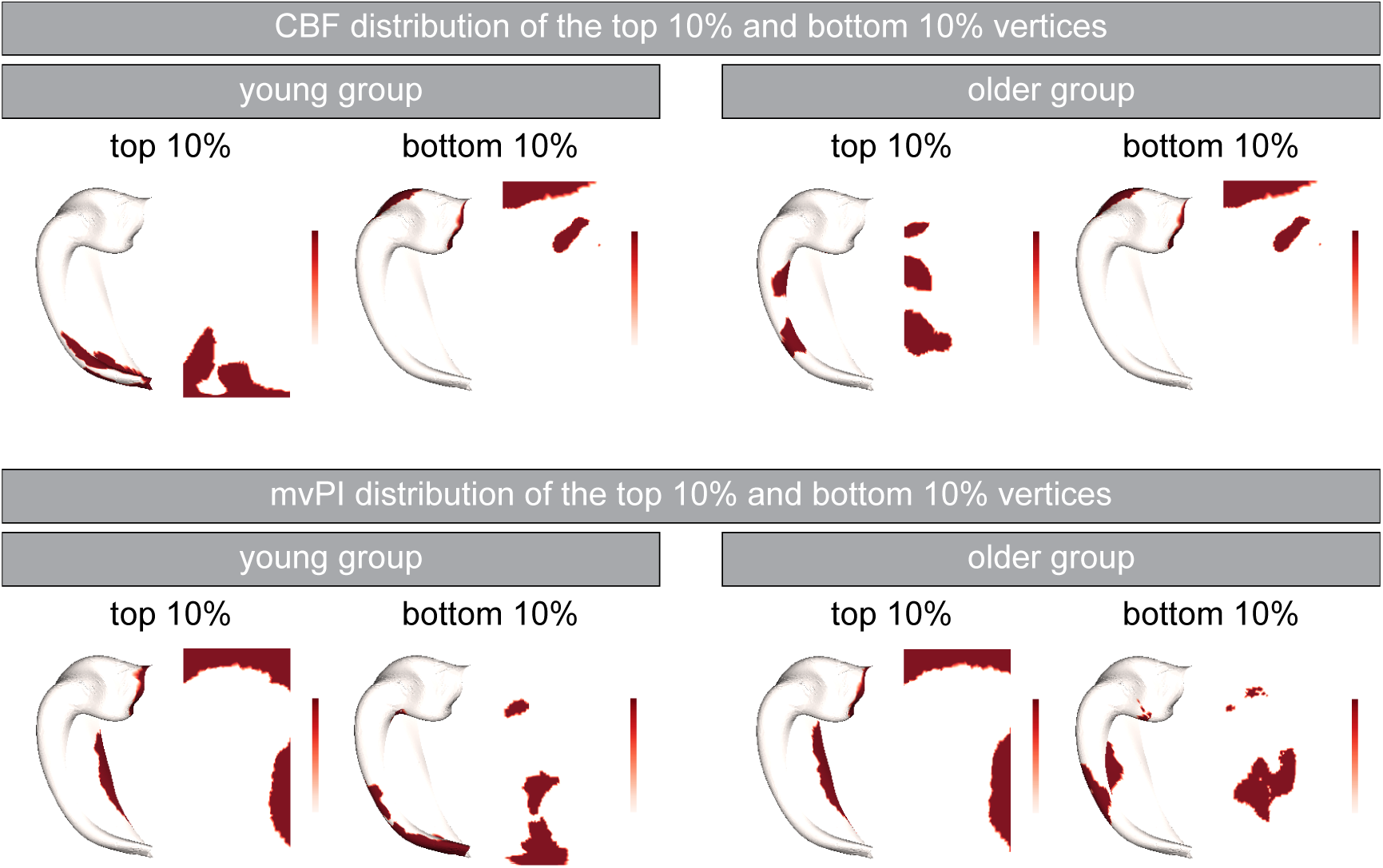
Spatial distribution of vertices with high and low CBF and mvPI in the hippocampus. Spatial distribution of vertices corresponding to the top and bottom 10% of CBF (top) and mvPI (bottom) values, shown separately for the young and older groups. For each vascular measure and age group, the selected vertices are displayed on both folded and unfolded hippocampal surfaces. These vertex sets were used for the frequency-resolved iEEG power comparisons shown in Fig. 4c**,d**.

**Fig. S4.**
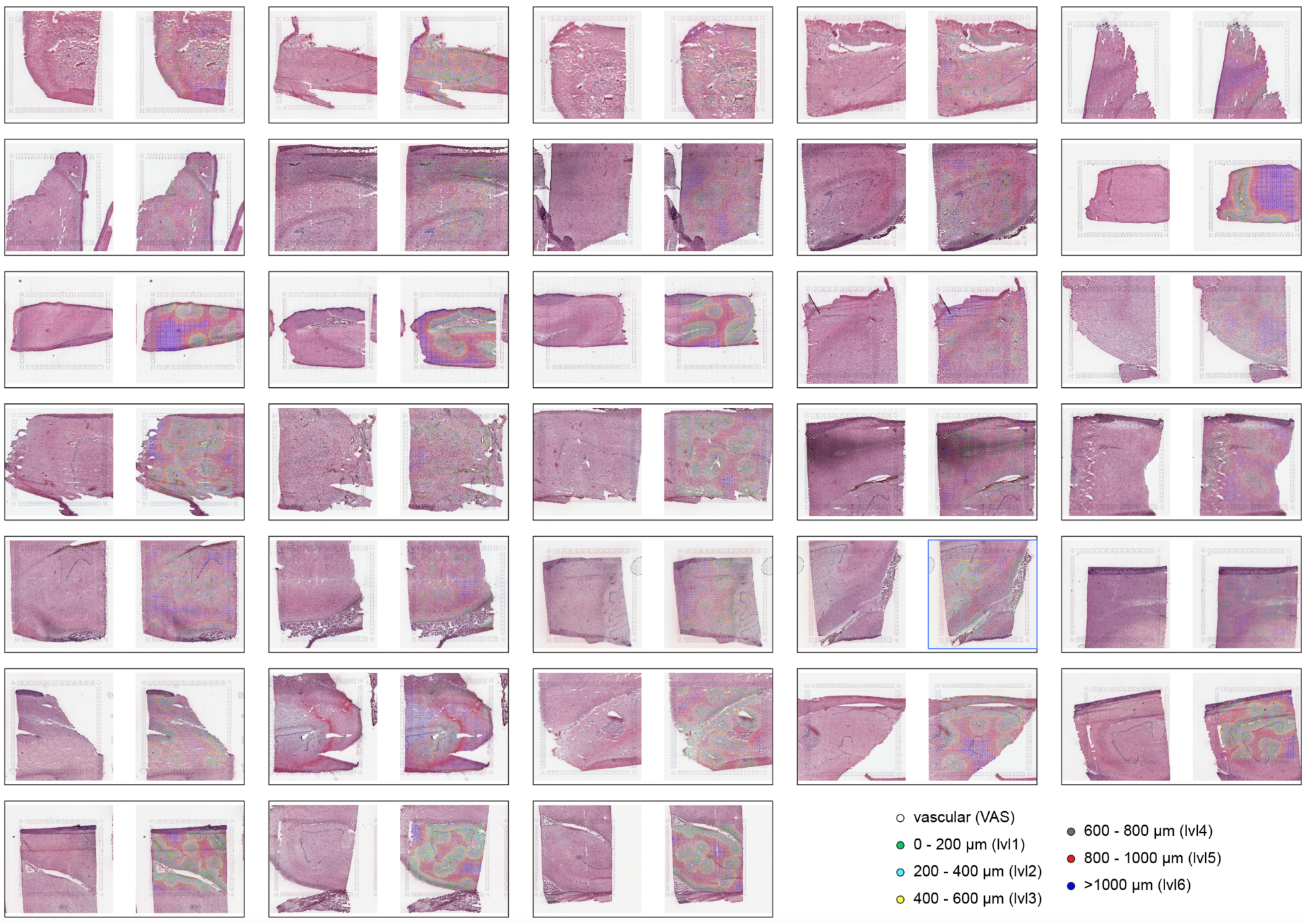
Histological images and vascular-distance gradients across all other spatial transcriptomic captures. H&E images of all 34 hippocampal spatial transcriptomic captures are shown together with the corresponding vascular-distance gradient (G_dist_) maps. For each capture, the original histological image is shown on the left and the G_dist_ classification overlaid on the tissue section on the right. Spots were classified as vascular (VAS) or into six progressively more distant bins according to their distance from the nearest delineated vessel: lvl1 (0–200 μm), lvl2 (200–400 μm), lvl3 (400–600 μm), lvl4 (600–800 μm), lvl5 (800–1000 μm), and lvl6 (>1000 μm).

**Fig. S5.**
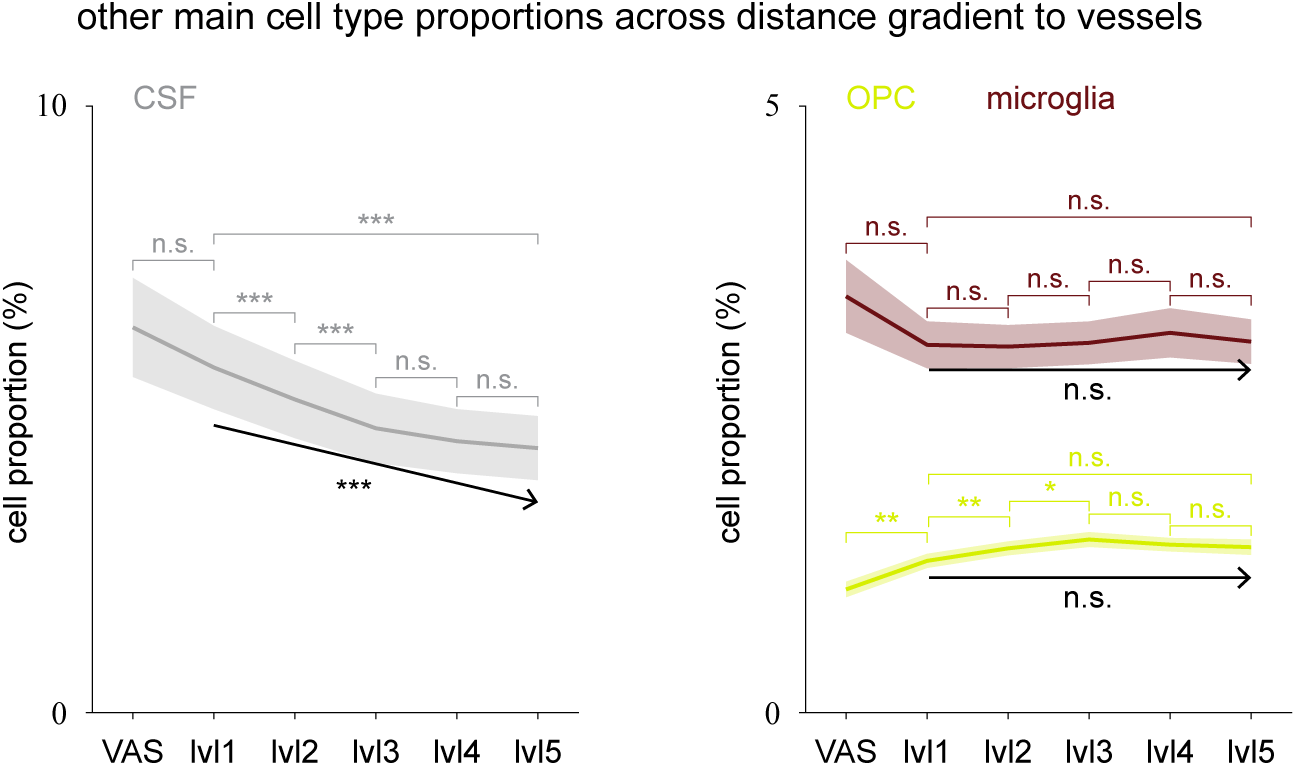
Other cell-type proportions across the vascular-distance gradient. RCTD-estimated proportions of CSF-associated cells, microglia and oligodendrocyte precursor cells (OPCs) across the vascular-distance gradient (Gdist), from vascular regions (VAS) to progressively more distant bins (lvl1– lvl5). Lines and shaded bands show the mean and SEM across captures. Horizontal brackets denote pairwise comparisons between distance bins (paired t-tests, BH-FDR corrected). p < 0.05, p < 0.01, p < 0.001; n.s., not significant. For each capture, a linear regression was fitted across lvl1–lvl5, and the resulting slope was tested against zero to assess the direction and significance of the within-capture distance-related trend.

**Fig. S6.**
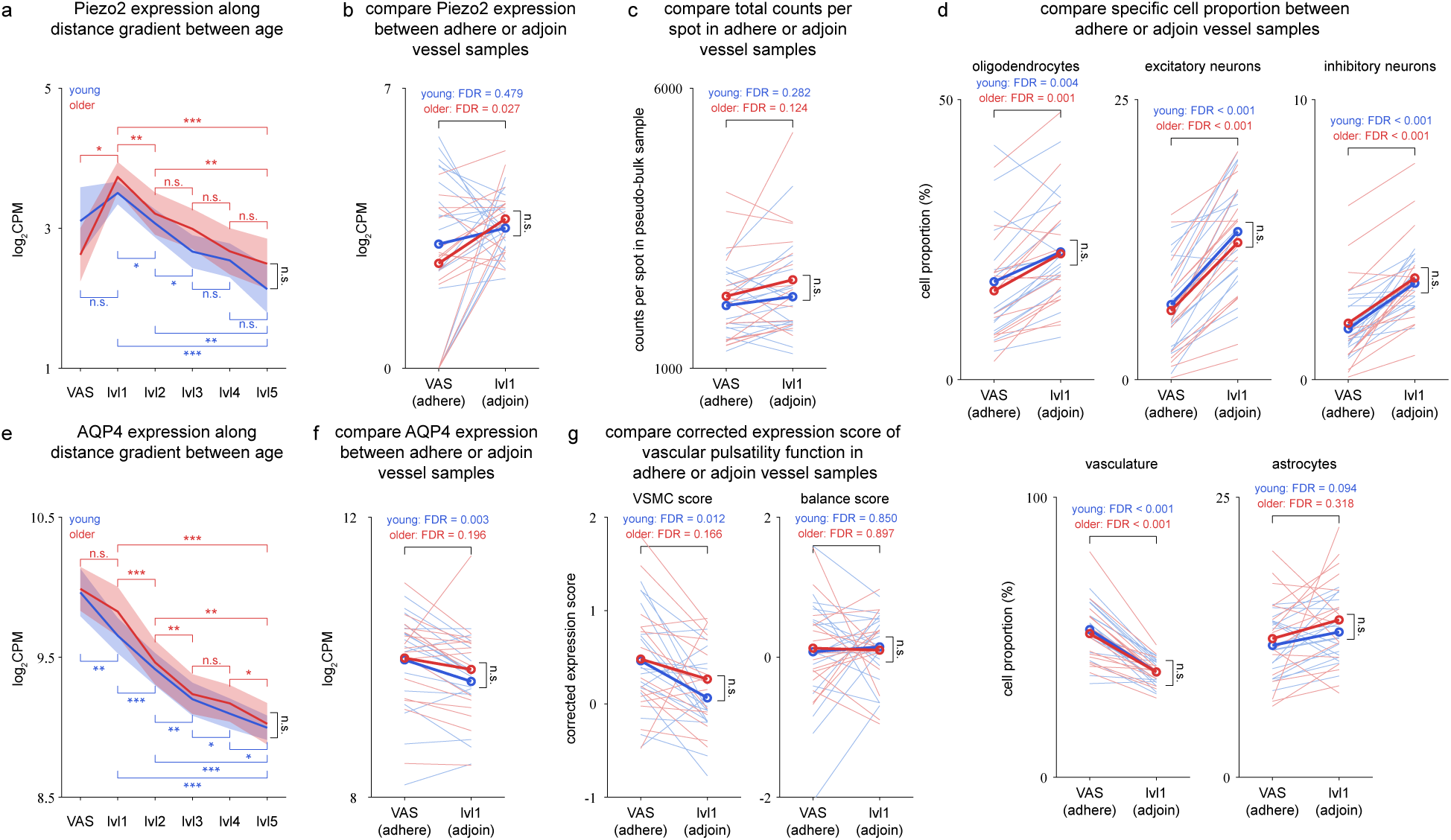
Distinct age-stratified patterns of perivascular PIEZO2 and AQP4 expression and vascular mechanical signatures. **a**, Piezo2 expression (log_2_CPM) across the distance gradient in young (blue) and older (red) adults. Lines and shaded bands give the mean and s.e.m. across captures. Colored bars denote within-age comparisons between bins (paired T-test, BH-FDR corrected). The black bar denotes the between-age comparison (unpaired t-test, BH-FDR corrected) and the age × G_dist_ interaction was not significant. **b**, Paired comparison of Piezo2 expression between vessel-adherent (VAS) and vessel-adjoining (lvl1) samples. Thin lines show individual captures, and thick lines show group means. Colored values give within-age paired comparisons (BH-FDR). The black bar denotes the between-age comparison (unpaired t-test, BH-FDR corrected) and the age × G_dist_ interaction was not significant. **c**, As in (**b**), for total counts per spot in the pseudo-bulk sample, confirming that the two sample types do not differ systematically in sequencing depth. **d**, As in (**b**), for the proportions of oligodendrocytes, excitatory neurons and inhibitory neurons (upper row), and of vasculature and astrocytes (lower right). **e-f**, As in (**a-b**), for AQP4. **g**, As in (**b**), for the corrected VSMC and compliance - stiffness balance scores defined in Fig. 5c**,d**.

**Fig. S7.**
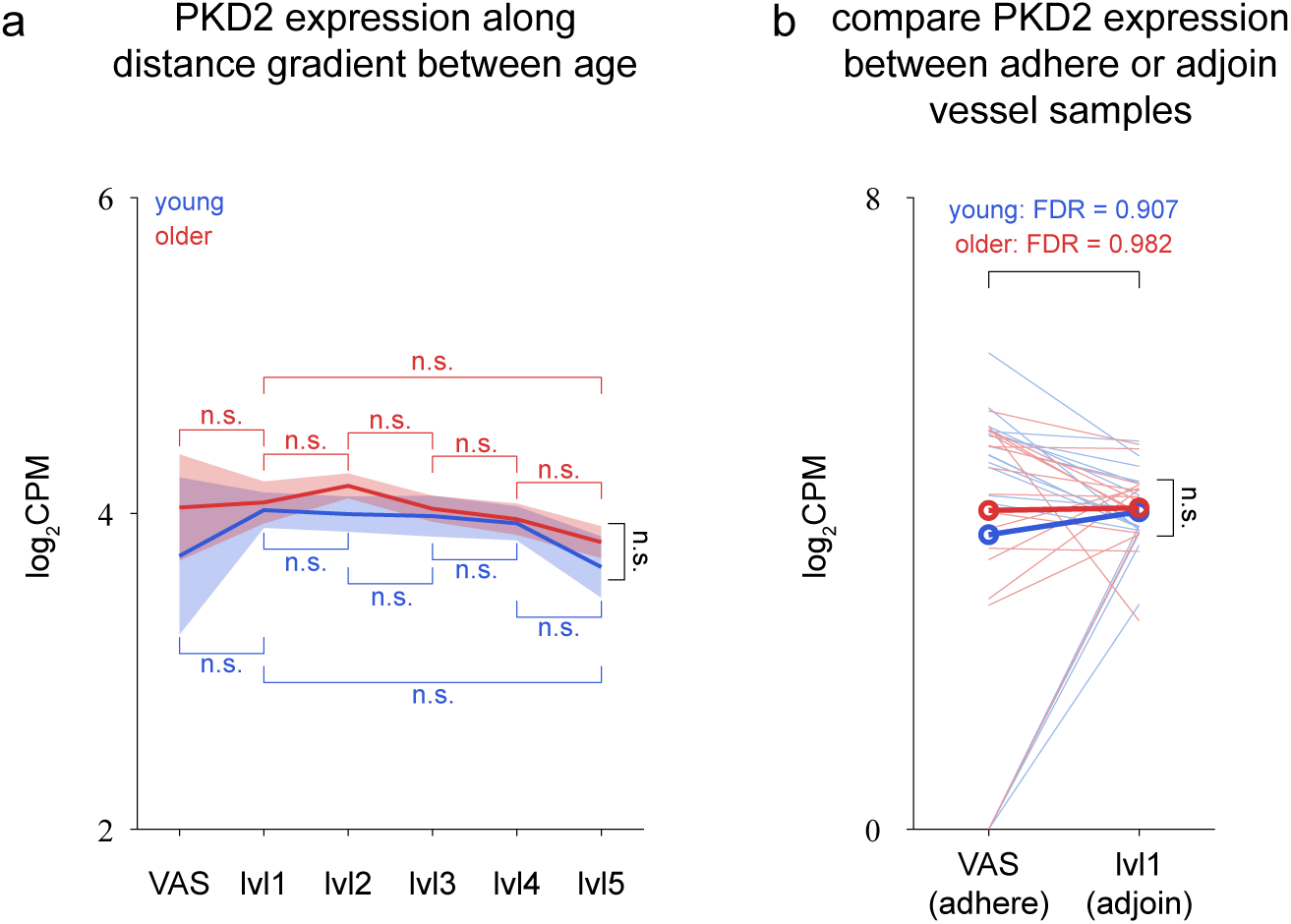
Age-stratified spatial expression patterns of PKD2 across G_dist_. Left, PKD2 expression (log2CPM) across the vascular-distance gradient in younger (blue) and older (red) hippocampal samples. Lines and shaded bands show the mean and s.e.m. across captures. Horizontal bars denote within-age pairwise comparisons between distance bins (paired t-tests, BH-FDR corrected). Right, paired comparison of PKD2 expression between vessel-adherent (VAS) and vessel-adjoining (lvl1) samples. Thin lines show individual captures, and thick lines show group means. FDR-corrected p values are shown for the within-age VAS–lvl1 comparisons. No significant VAS–lvl1 difference was detected in either age group. n.s., not significant.

## Notes

### Competing Interest Statement

The authors have declared no competing interest.

